# Evolution is in the details: Regulatory differences in modern human and Neanderthal

**DOI:** 10.1101/2020.09.04.282749

**Authors:** Harlan R. Barker, Seppo Parkkila, Martti E.E. Tolvanen

## Abstract

Transcription factor proteins play a critical role in the regulation of eukaryote gene expression via sequence-specific binding to genomic locations known as transcription factor binding sites.

We studied sites of genomic variation between modern human and Neanderthal promoters. We detected significant differences in the binding affinities of 110 transcription factors to the promoters of 74 target genes. The transcription factors were enriched for terms related to vision, motor neurons, homeobox, and brain, whereas the target genes and their direct interactors were enriched in terms related to autism, brain, connective tissue, trachea, prostate, skull morphology, and vision. Secondary analysis of single-cell data revealed that a subset of the identified transcription factors (CUX1, CUX2, ESRRG, FOXP1, FOXP2, MEF2C, POU6F2, PRRX1 and RORA) co-occur as marker genes in L4 glutamatergic neurons. The majority of these genes have known roles in autism and/or schizophrenia and are associated with human accelerated regions (elevated divergence in humans vs. other primates). These results support the value of gene regulation studies for the evolution of human cognitive abilities and the neuropsychiatric disorders that accompany it.

## Introduction

### North of Eden

Humans and their hominin relatives have been leaving Africa in waves for the past two million plus years. Various environmental and cultural pressures have impacted each diaspora in different ways, subsequently producing adaptations reflected in physiology, immunity, and brain size. Recently, the discovery and sequencing of DNA from the remains of Neanderthal [1–4] and Denisovan [5,6] have allowed direct comparisons of DNA from modern and ancient hominids. Among the observed genomic variations between modern humans and Neanderthals, a limited number have been identified in gene coding regions. Some of these genes are known to affect cognition and morphology (cranium, rib, dentition, and shoulder joint) [1], pigmentation and behavioral traits [2], and brain development [7]. However, as has been noted before, there is a paucity of coding variations to explain the differences between related species; the genome of the Altai Neanderthal reveals just 96 fixed amino acid substitutions, occurring in 87 proteins [7]. Unsurprisingly, a much larger set of variants are observed in intergenic regions, owing not only to the fact that these are comparatively much larger regions but also to the expectation of less conservation in what until recently was often termed "junk DNA". While variants in coding regions can directly affect protein structure, those found in intergenic regions may affect the regulation of gene expression through alternative binding of transcription factors in promoters and enhancers and the expression of noncoding RNAs. What may surprise some is the cumulative effect of numerous small — and large — changes to gene expression arising owing to these manifold intergenic changes, which may ultimately serve as the engine of speciation. Indeed, introgressed Neanderthal DNA is more depleted in regulatory regions than in those encoding proteins [8]. Using a new computational tool that incorporates numerous transcription-relevant genomic features, we sought to reveal how the comparative differences in these regions may affect regulatory differences between modern humans and Neanderthals. Specifically, our aim was to identify those gene-regulating transcription factors whose binding to DNA may vary between these species of hominids and thus drive the differences between them.

## Methods

### Analysis of modern human vs. Neanderthal genetic variation

The Neanderthal Genome Project has cataloged a total of 388,388 SNPs in modern human and Neanderthal genomes [1]. These were further reduced to 21,990 SNPs in the proximal promoters (-2,500 to +2,500 nts relative to the TSS) of those transcripts, which are defined by Ensembl as "protein-coding" (Figure 1). Using TFBSFootprinter [9–12], a tool we have developed for the prediction of TFBSs, a 50 bp region centered on each SNP was analyzed for the binding of 575 TFs for both the modern human version and the Neanderthal variant. TFBSFootprinter automatically retrieves the human sequences at a target region, and custom Python scripts were used to modify these sequences for the Neanderthal variant. All TFBS predictions that overlapped with the target SNP position were retained. The complete result set was then reduced on the basis of the combined affinity score p value via a Benjamini‒Hochberg-derived critical p value corresponding to a false discovery rate (FDR) cutoff of 0.01 to address multiple testing. For each putative TFBS meeting the cutoff in either subspecies, the corresponding matched pair of PWM scores was retained. To identify significantly different scoring for TFs between subspecies, for each TF, via the compiled matched scores, the Wilcoxon rank test was performed via the SciPy stats Python library [13], and subsequent results were filtered via a Benjamini‒Hochberg-derived critical p value corresponding to an FDR cutoff of 0.10 to address multiple testing.

**Figure 1.**
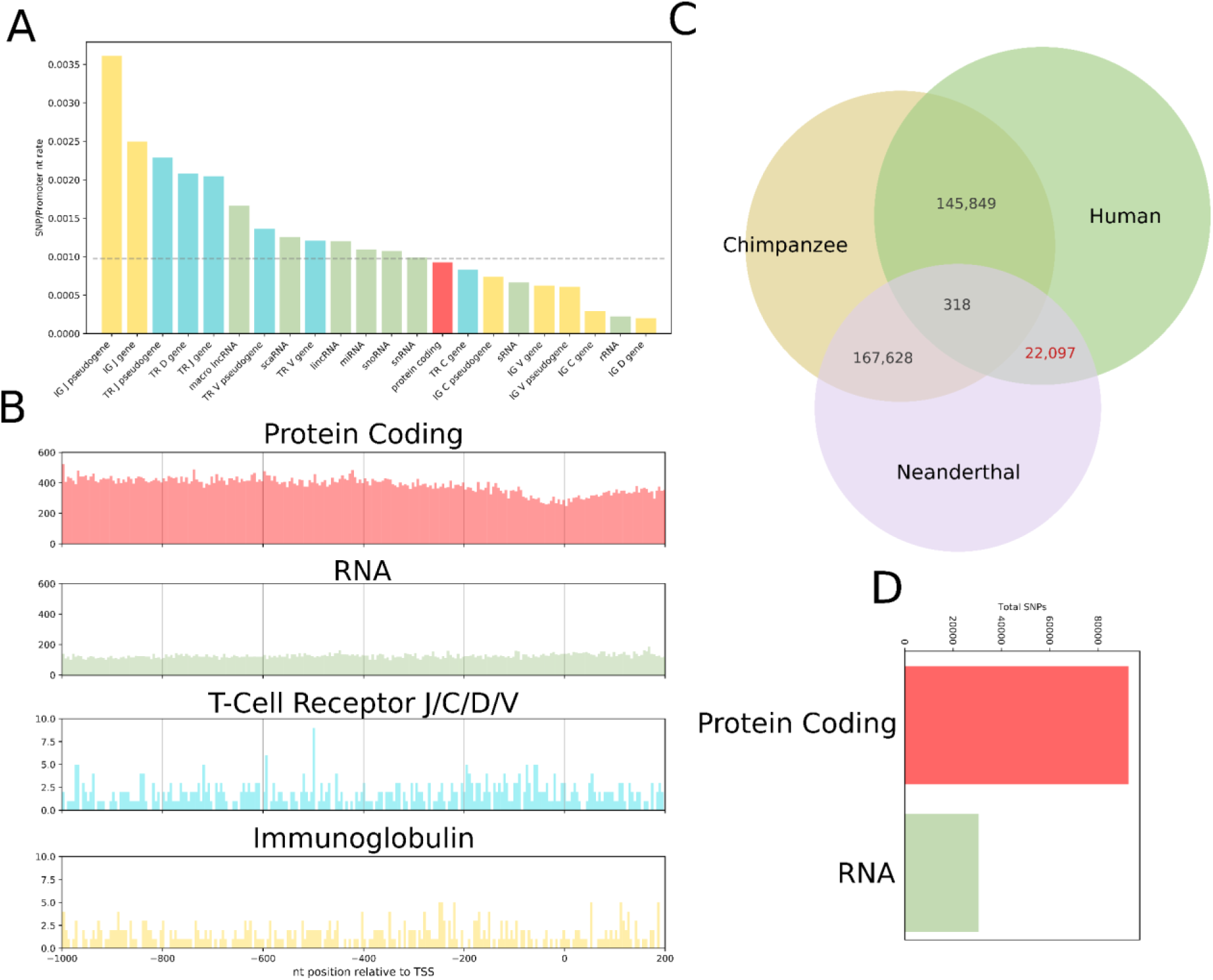
Modern human and Neanderthal variants used in the analysis. (**A**) Incidence rate of SNPs/nt in human promoters of the various transcript classes, as defined by Ensembl. The bars are colored by transcript class and correspond with those in panel B. (**B**) Number of observed modern human/Neanderthal SNPs at each nt location in the sets of promoters of various transcript types. A visible depletion of SNPs is observed for the promoters of protein-coding transcripts corresponding with increasing proximity to the TSS (nt position 0). (**C**) Counts of total SNPs in promoters of protein-coding transcripts between the three species cataloged in the Neanderthal Genome Project. (**D**) Comparison of counts of human vs. chimp SNPs in promoters of protein-coding vs RNA transcripts. IG J, immunoglobulin joining chain; TR J, T-cell receptor joining chain; TR D, T-cell receptor diversity chain; TR V, T-cell receptor variable chain; IG C, immunoglobulin constant chain; IG V, immunoglobulin variable chain; IG D, immunoglobulin diversity chain.

A total of 108 TF models were identified as scoring differently across human vs. Neanderthal SNP locations. For each of these genes, we extracted RNA-Seq data from the FANTOM dataset (across all CAGE peaks associated with that TF) and retained the data for the 100 tissues with the highest aggregate expression across all of the target TF genes. In the case of heterodimer JASPAR TF models (e.g., FOS::JUN, NR1H3::RXRA, and POU5F1::SOX2), the expression of each TF component gene was used. The expression data were extracted as TPM values and normalized by log_2_ transformation. From the subsequent normalized expression data values, hierarchical clustering was performed and visualized via SciPy, Matplotlib, and Seaborn Python libraries [13–15].

The results of hierarchical clustering revealed a cluster of TF genes that were uniquely expressed in neural and immune tissues. These gene sets were used to perform PANTHER-based gene ontology enrichment analysis (using the www.geneontology.org web server) [16,17], with the default statistical settings using Fisher’s exact test method and an FDR threshold of p<0.05 (default setting).

### Enrichment analysis of DB TFs and their target genes

The OmniPath DB compiles protein annotation data from several tens of publicly available datasets. The complete set of annotation data was downloaded and filtered to annotations from PanglaoDB and UniProt tissue, family, and keyword classifications. The set of 110 DB TFs was analyzed for enrichment for all annotation labels via Fisher’s exact test and compared with the full set of 529 human JASPAR TF genes, with Benjamini‒Hochberg correction for multiple testing (FDR cutoff value of 0.1).

The OmniPath DB has likewise compiled protein interactions (human, mouse and rat) from many publicly available datasets. The complete set of interactions was downloaded and filtered to determine posttranslational human protein‒protein interactions. A total of 805 interactors were identified for the set of 74 DB TF target genes, and the combined list (879 genes) was analyzed for gene ontology enrichment analysis via the geneontology.org webserver [16,17], with the default settings of Fisher’s exact test and an FDR cutoff of 0.05 for multiple testing correction.

### Analysis of the brain expression of differentially binding TFs

Expression data in the form of reads per kilobase per million reads (RPKM) were extracted for 26 unique tissues across 31 timepoints (from 8 weeks postconception to 40 years after birth) from the Allen Brain Atlas (brainspan.org). The RPKM values were then converted to TPM values and log_2_ transformed. Ages were grouped into three phases of growth for simplicity of analysis and interpretation: pcw (8–37 weeks post conception), early (4 months–4 years), and late (8 years–40 years). Corresponding to these age groups, log-transformed TPM values for each tissue were grouped and averaged and used to perform cluster map analysis to identify grouping tissues at time points with similar expression profiles.

### Analysis of brain samples via scRNA-Seq

The Allen Brain Atlas has performed single-nucleus RNA-Seq analysis of 49,417 nuclei derived from 8 brain cortex regions within the middle temporal gyrus (MTG), anterior cingulate gyrus (CgG), primary visual cortex (V1C), primary motor cortex (M1C), primary somatosensory cortex (S1C) and primary auditory cortex (A1C) [18]. These data were downloaded as a count matrix along with a table of the associated metadata (https://portal.brain-map.org/atlases-and-data/rnaseq/human-multiple-cortical-areas-smart-seq) and loaded into the Python library SCANPY [19]. Using a modified workflow described previously [20], samples were filtered by Gaussian fit of the read count (300,000<x<3,500,000), expressed gene count (2,000<x), and number of cells in which a gene was expressed (>50), resulting in a final count of 46,959 cells and 42,185 genes for further analysis.

Using SCANPY, counts were normalized by cell (‘pp.normalize_total’; target_sum=1,000,000), log transformed (‘pp.log1p’), highly variable genes identified (‘pp.highly_variable_genes’; flavor=‘seurat_v3’; n_top_genes=5,000; layer=’counts’), principal component analysis (‘pp.pca’; n_comps = 15; svd_solver=’arpack’), and k-nearest neighbors (‘pp.neighbors’; n_neighbors=15). The expression relationships between cells were graphically visualized with the Python implementation [21] of the ForceAtlas2 [22] graph layout algorithm, as called in Scapny (‘tl.draw_graph’ and ‘pl.draw_graph’).

Annotation data regarding brain region, cortical layer, and GABAergic/glutamatergic/nonneuronal cell type features were extracted from Allen Brain Atlas sample data for mapping onto derived cell type clusters. The top 100 marker genes for each cell type+cortical layer (CT/CL) cluster (e.g., excitatory L4) were identified as those with higher expression unique to each cluster by the Welch t test in SCANPY (‘tl.rank_genes_groups’). The expression of the DB TF list genes, which were identified as marker genes in a CT/CL cluster, was mapped onto cluster figures.

Gene ontology analysis of target cluster marker genes was performed via the Protein Analysis Through Evolutionary Relationships (PANTHER) tool at the Geneontology.org web server [17]. Ontological terms associated with overabundance among cluster marker gene lists were established via Fisher’s exact test, and the results were filtered by an FDR<0.05 (default setting); analyses were performed for biological process, molecular function, and cellular component terms. Disease term gene ontology analysis was performed via Enrichr [23,24], which is based on the ontology compiled by DisGeNET [25].

## Results

### High-scoring TFBSs differ between modern humans and Neanderthals and are enriched in homeobox and the brain

In the analysis of 21,990 SNPs identified in the comparison of the modern human and Neanderthal promoteromes—the collection of all human/Neanderthal proximal promoters of protein-coding genes—a total of 108 TF models, representing 110 unique differentially binding (DB) TF proteins—showed a significant difference in scoring (Wilcoxon rank statistical test) between the two hominin species (Table 1). Compared with the complete set of JASPAR TF genes from which they are derived, DB TF genes are strongly enriched among the homeobox genes (78/110; Fisher’s exact test: odds ratio [OR] 12.40; p value 2.96 × 10^-17^) and enriched in brain-specific expression (79/110; OR 2.86, p value 3.48 × 10^-6^). Similarly, compared with all protein-coding genes, the complete set of TFs cataloged in the JASPAR database are themselves very strongly enriched for homeobox genes (OR 85.16; p value 5.79 × 10^-177^) and moderately enriched for brain, eye, and retina genes (OR 1.80; p value 3.50 × 10^-11^). DB TF genes were underenriched among immune-specific genes (29/110) compared with the JASPAR set (OR 0.37; p value 1.44 × 10^-5^).

**Table 1.**
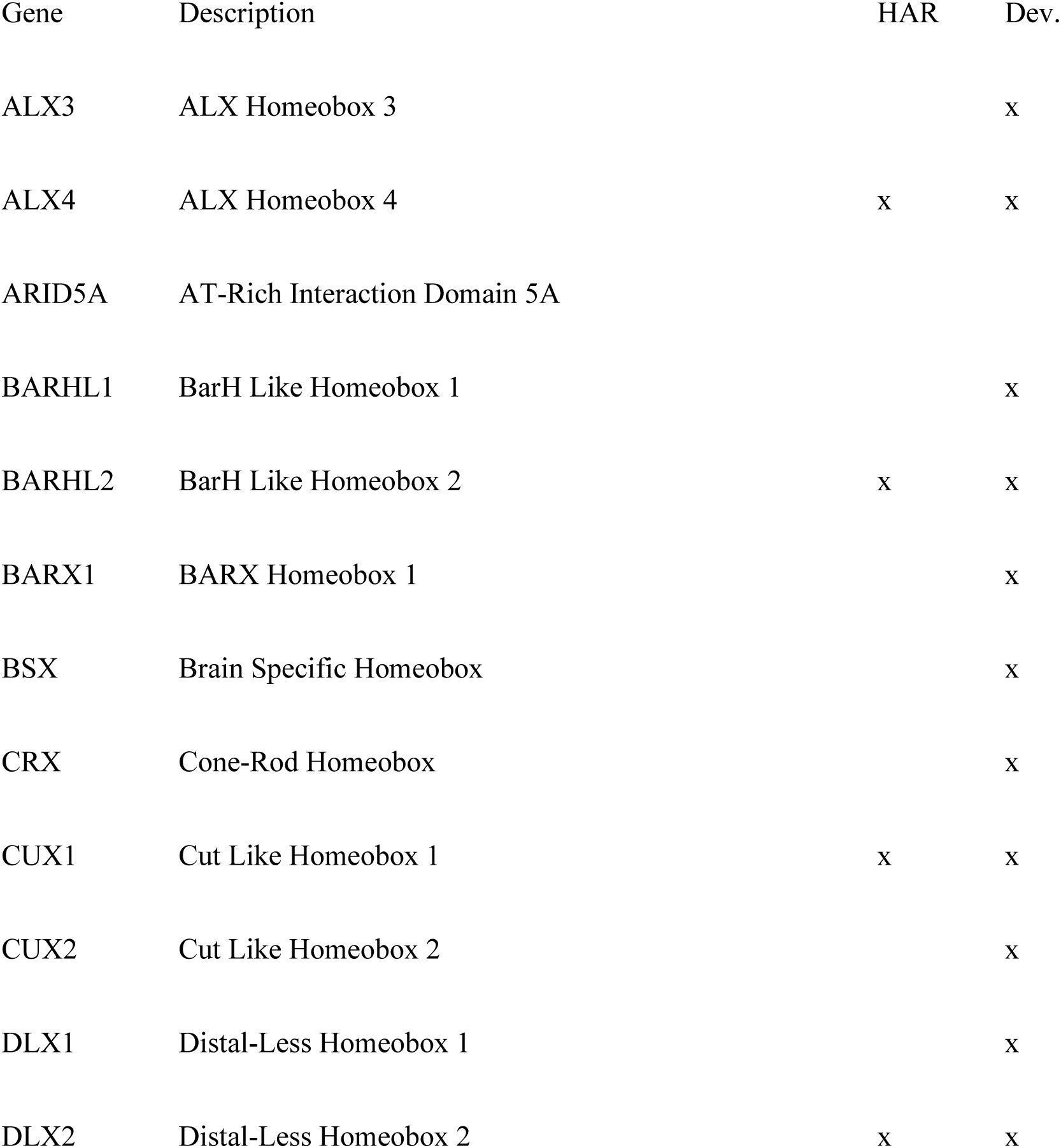
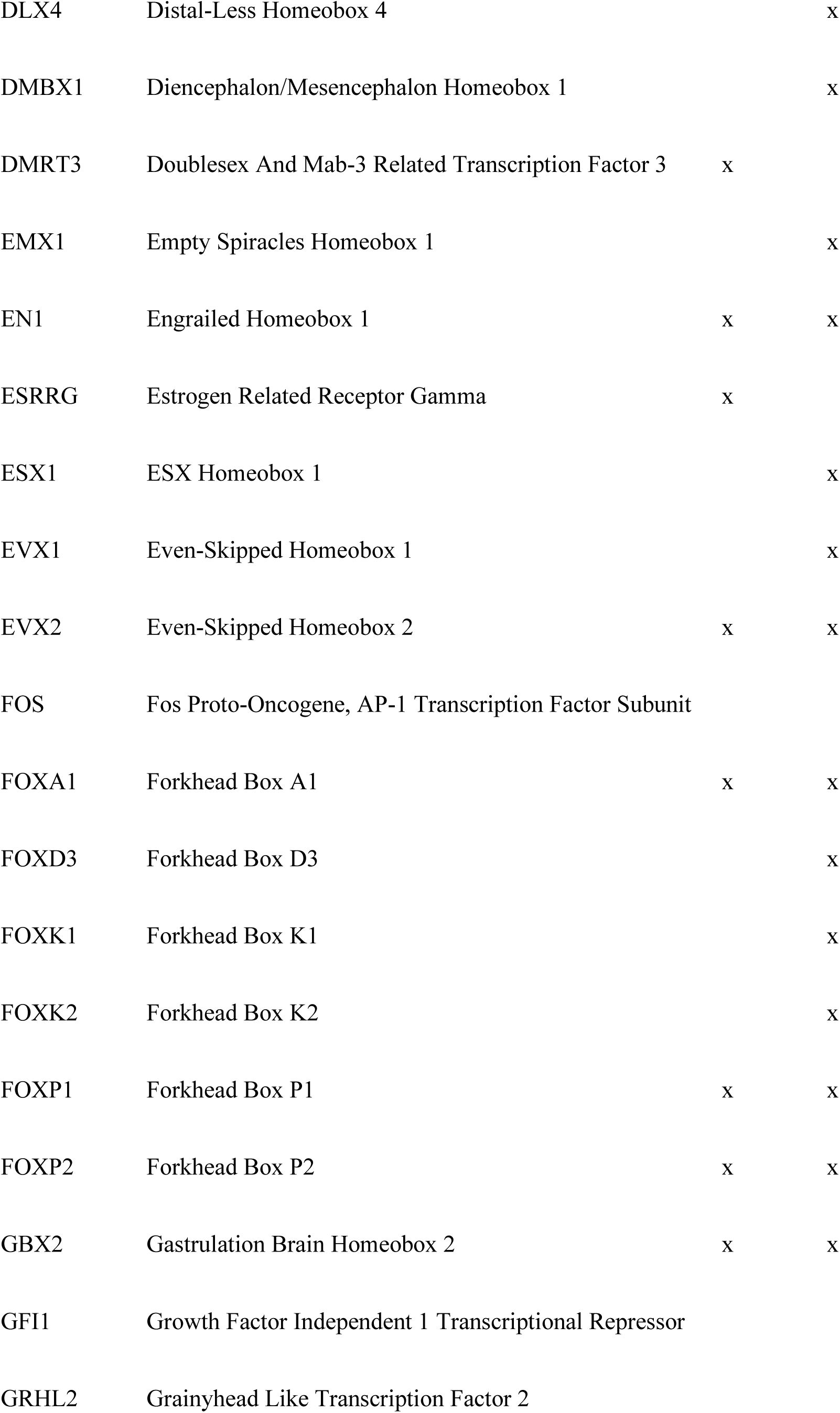
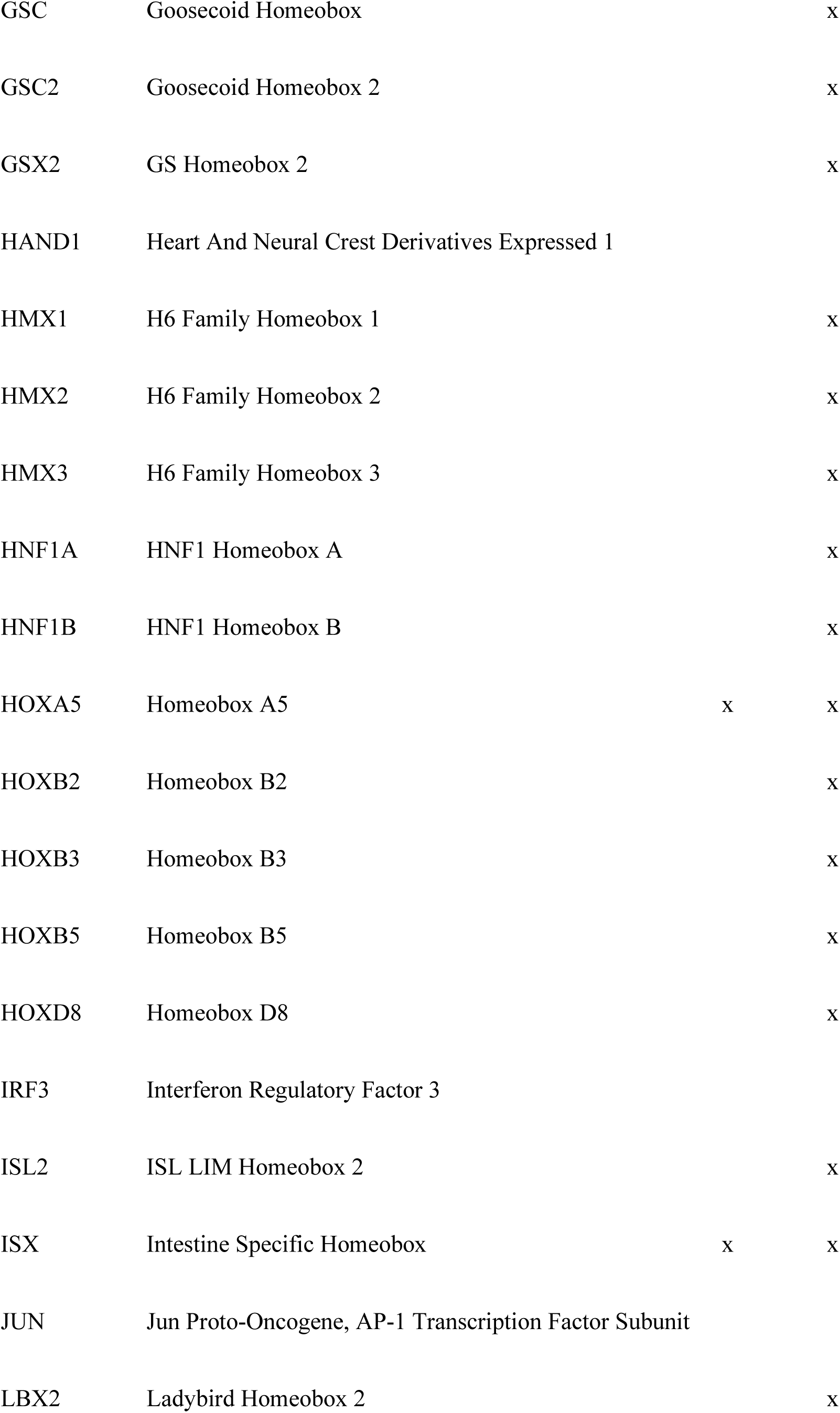
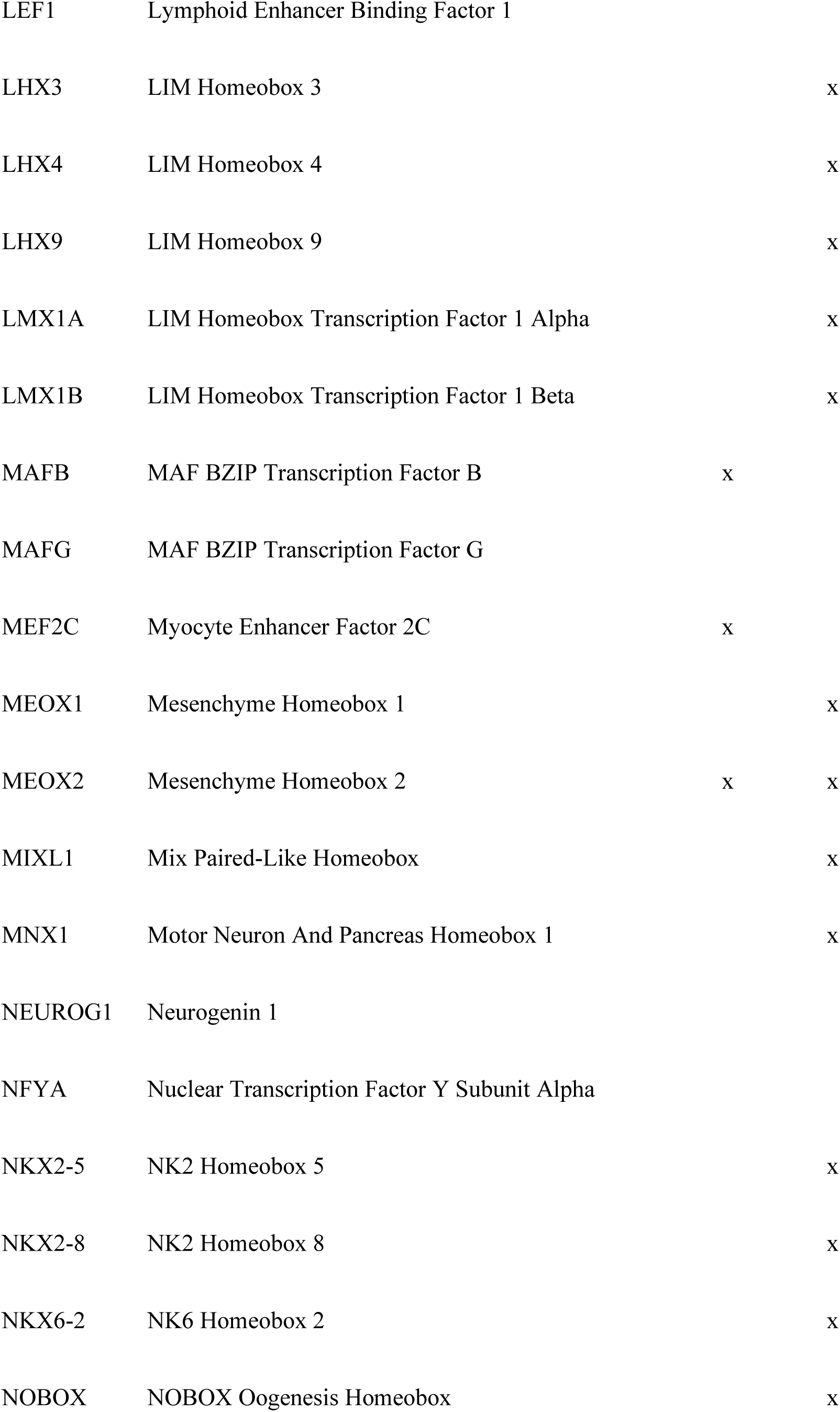
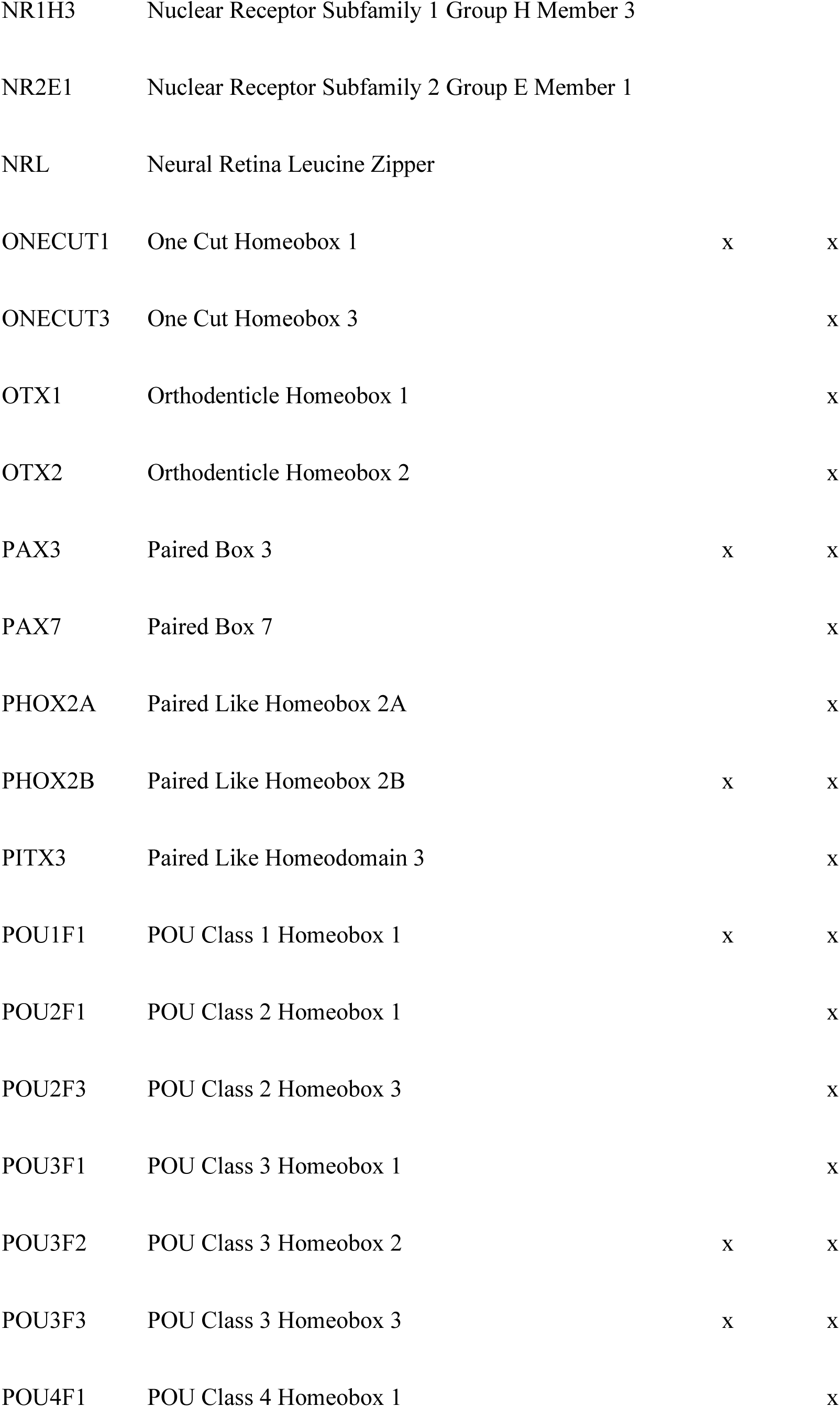
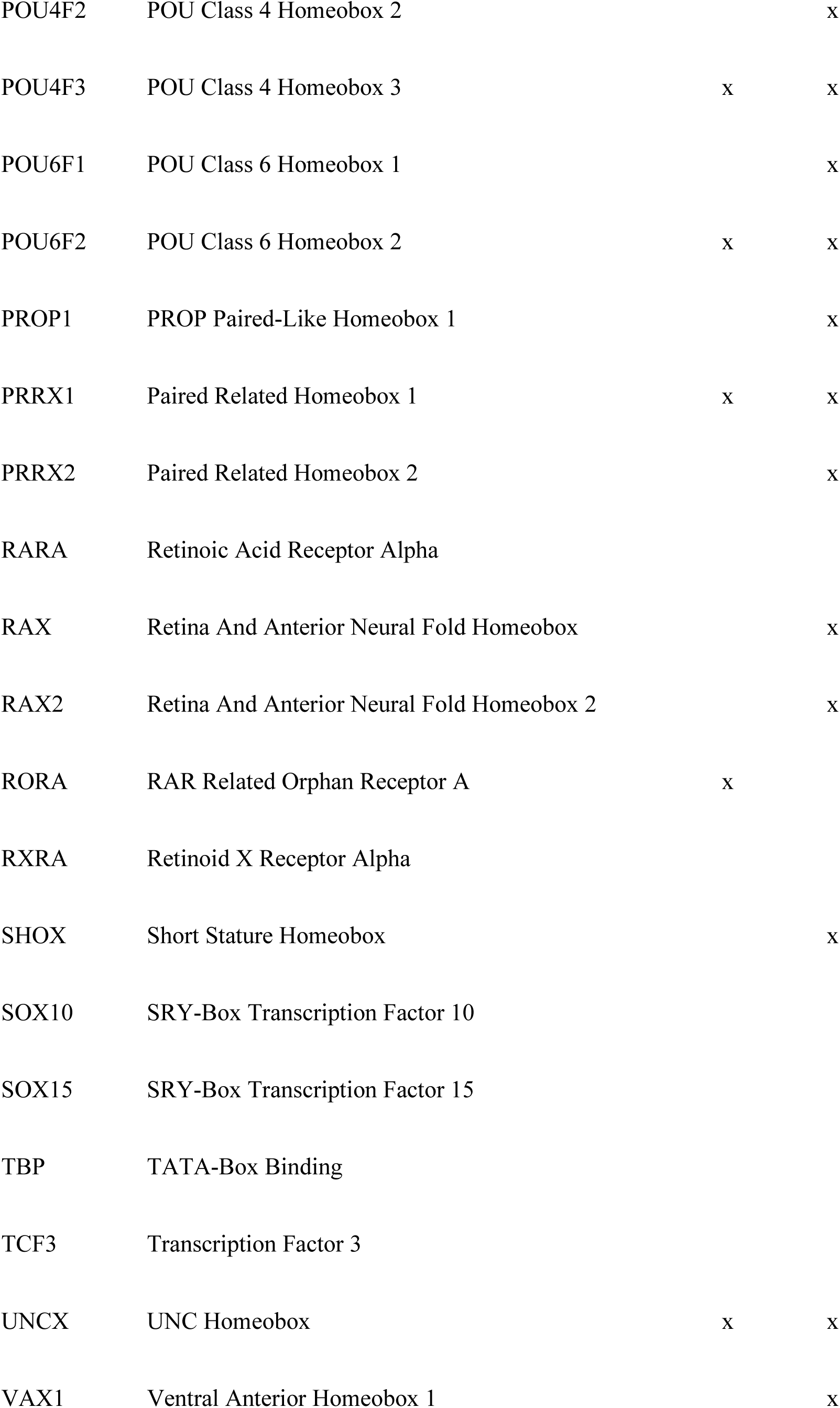
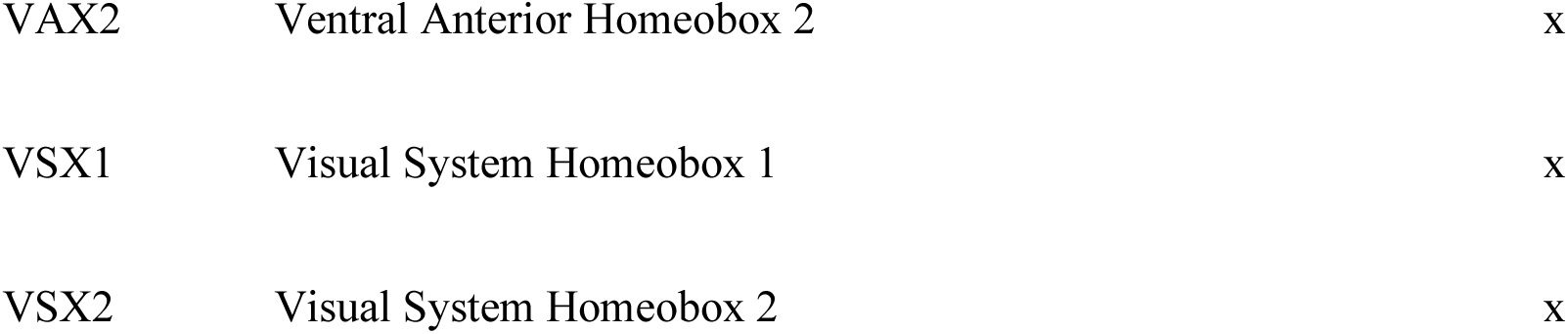
TFs with differential binding affinities in modern humans vs. Neanderthal proximal and semi-distal promoter regions. Dev., development-associated gene; HAR, human accelerated region.

### DB TFs enriched for vision, brain, and developmental terms

Enrichment analysis of the DB TF genes compared with the JASPAR TF genes was performed via the OmniPath annotation database [26,27] via Fisher’s exact test, with Benjamini‒Hochberg correction for multiple testing and an FDR of 0.1. The DB TFs were over-enriched for ‘Retina’ (OR *inf*; FDR 0.015), ‘Motor neurons’ (OR 28.4; FDR 0.013), ‘CUT homeobox’ (OR 15.8; FDR 0.042), ‘Vision’ (OR 9.93; FDR 0.058), ‘Sensory transduction’ (OR 9.93; FDR 0.058), ‘Bicoid’ (OR 9.42; FDR 0.0078), ‘Dwarfism’ (OR 4.61; FDR 0.058), ‘Developmental protein’ (OR 2.89; FDR 1.56 × 10^-4^), ‘Brain’ (OR 2.26; FDR 0.045), and ‘Ectoderm’ (OR 2.14; FDR 0.016), with the full results available in Supplementary Table 1.

### DB TF target genes

A total of 74 genes were identified, hereafter referred to as ‘DB TF target genes’, whose promoters contain MH vs. Neanderthal SNPs where differential binding occurs by our determined set of statistically significant DB TFs (Table 2). In all of the DB TF target genes, at least one MH vs. Neanderthal SNP position has been identified as an eQTL affecting the expression of the target gene in whose promoter it occurs, as defined by data from the GTEx database (data not shown).

**Table 2.**
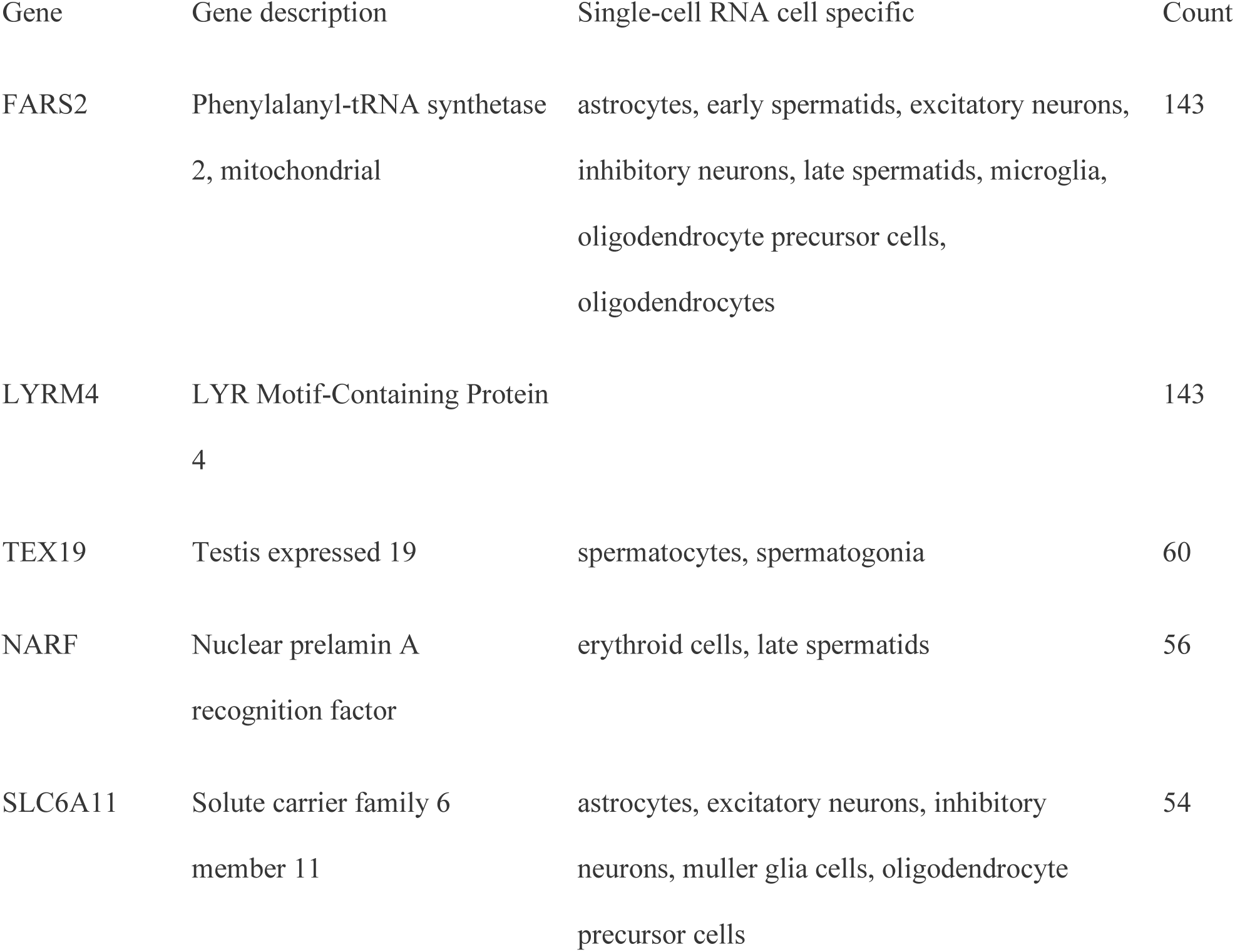
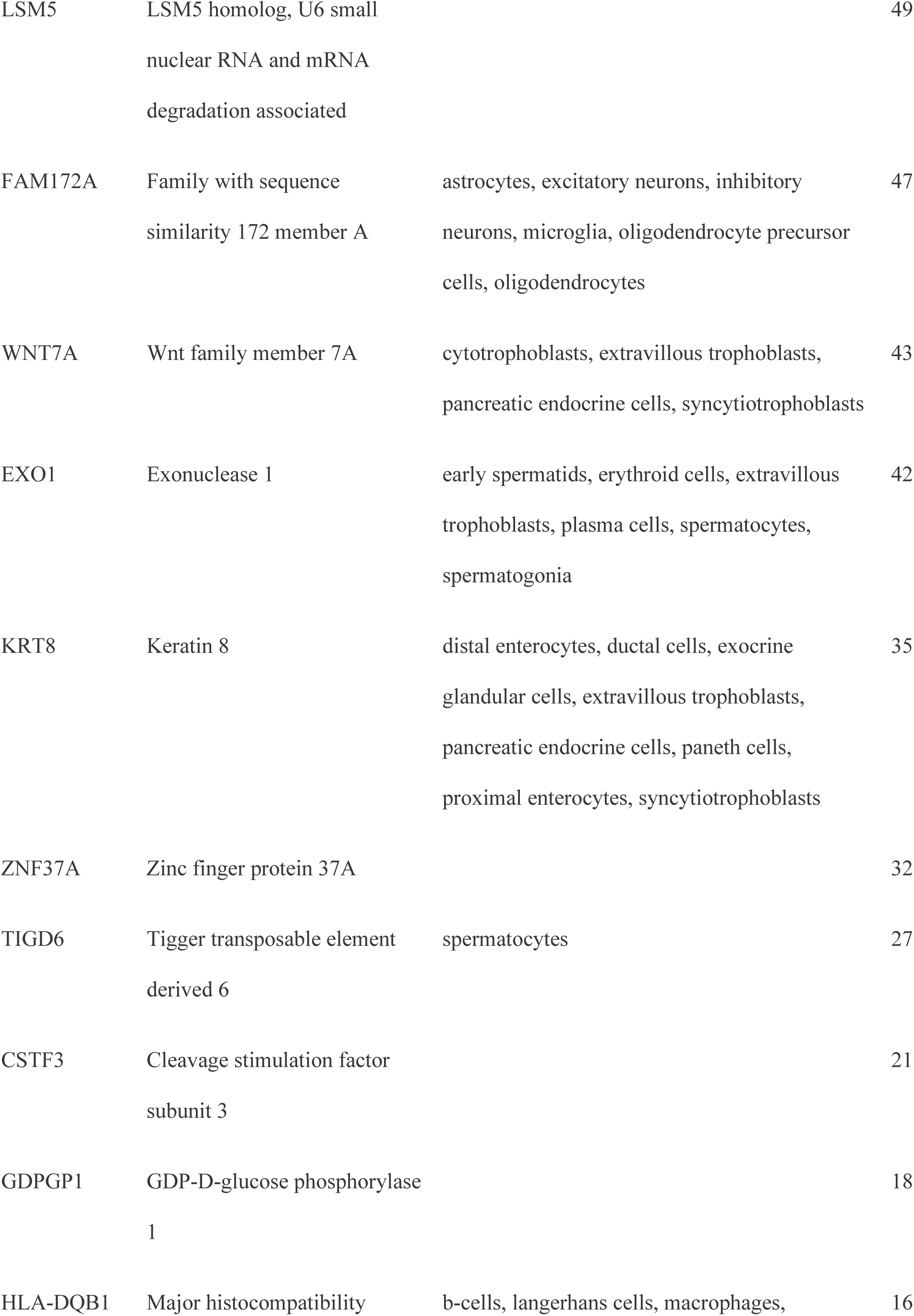
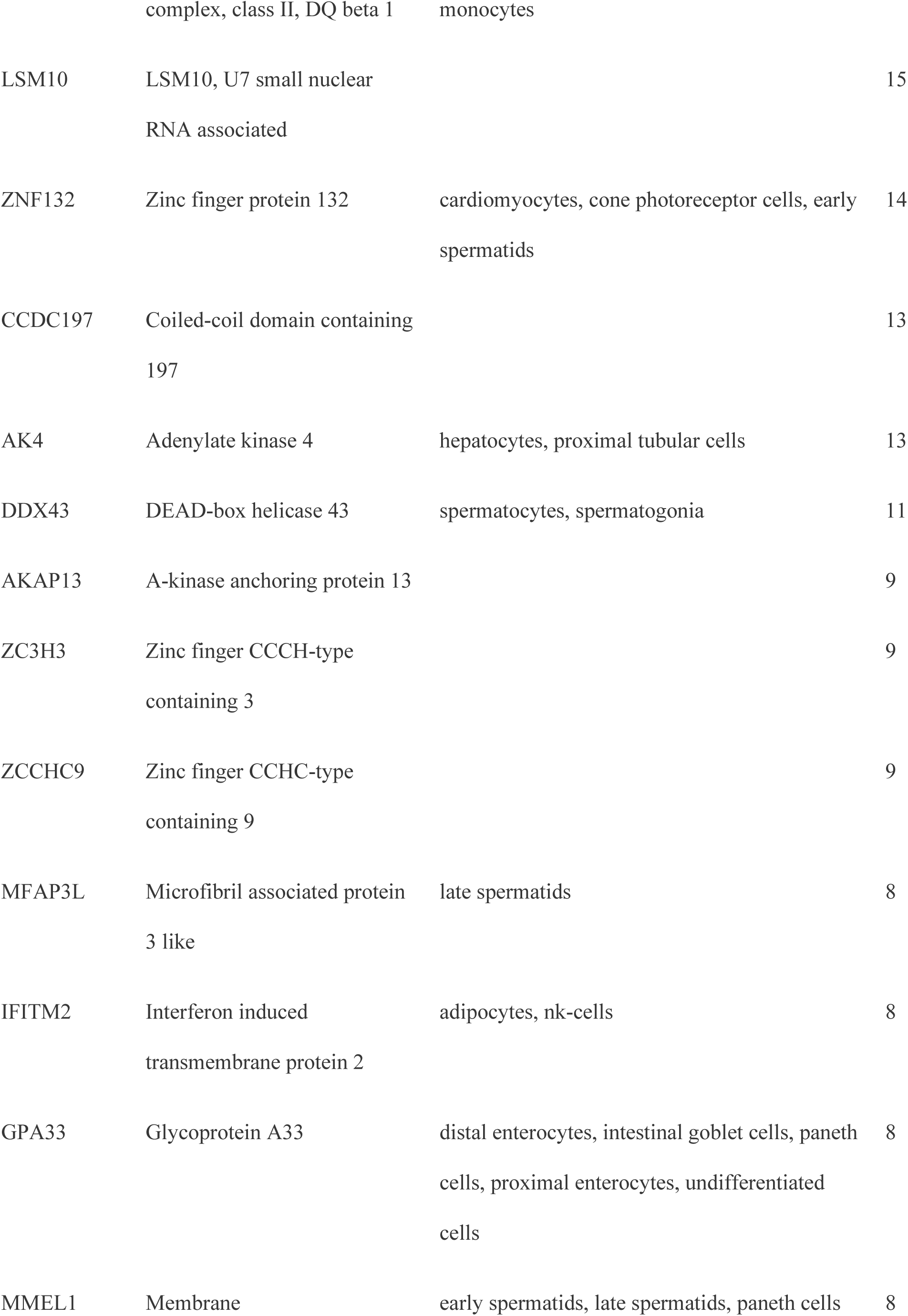
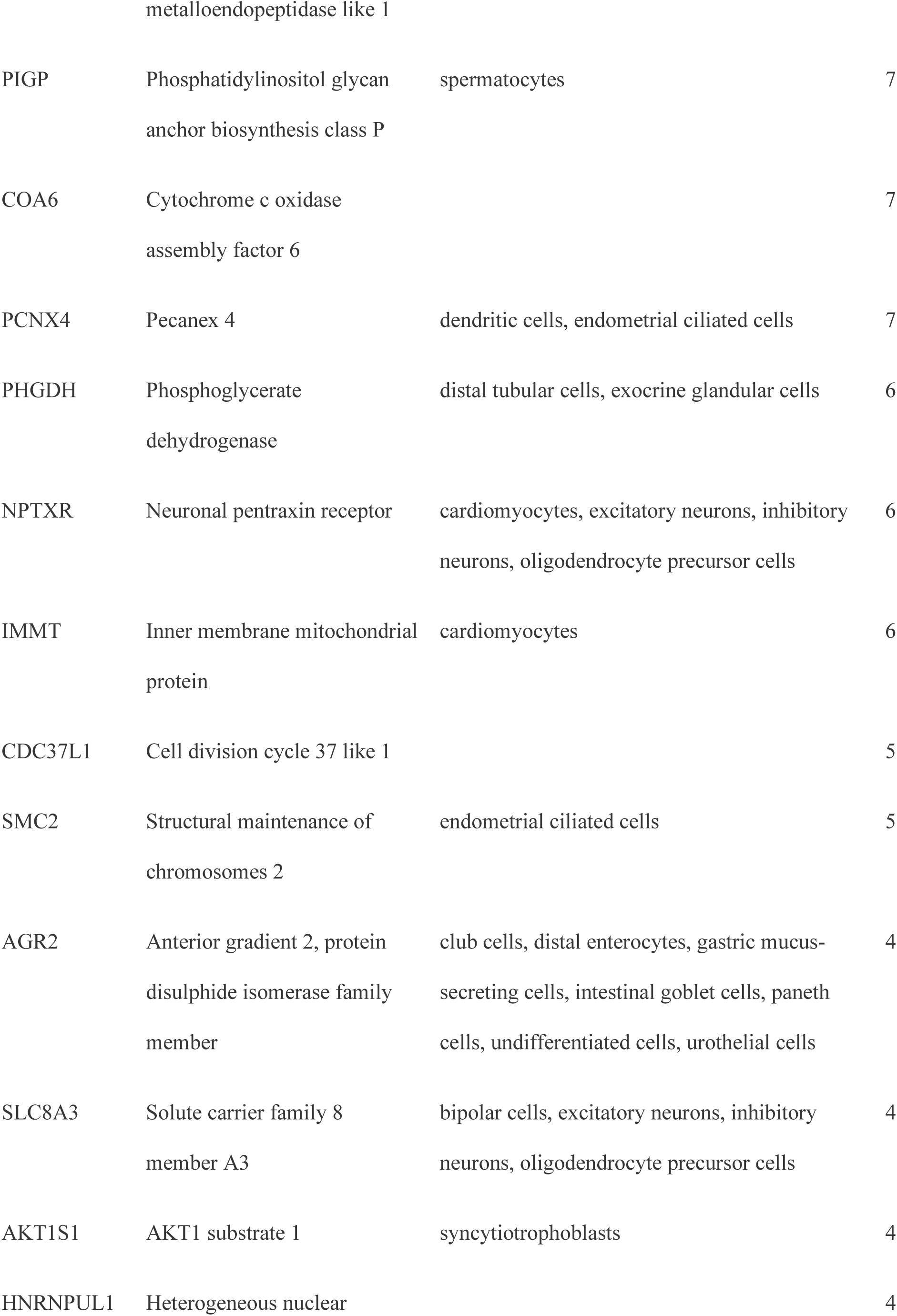
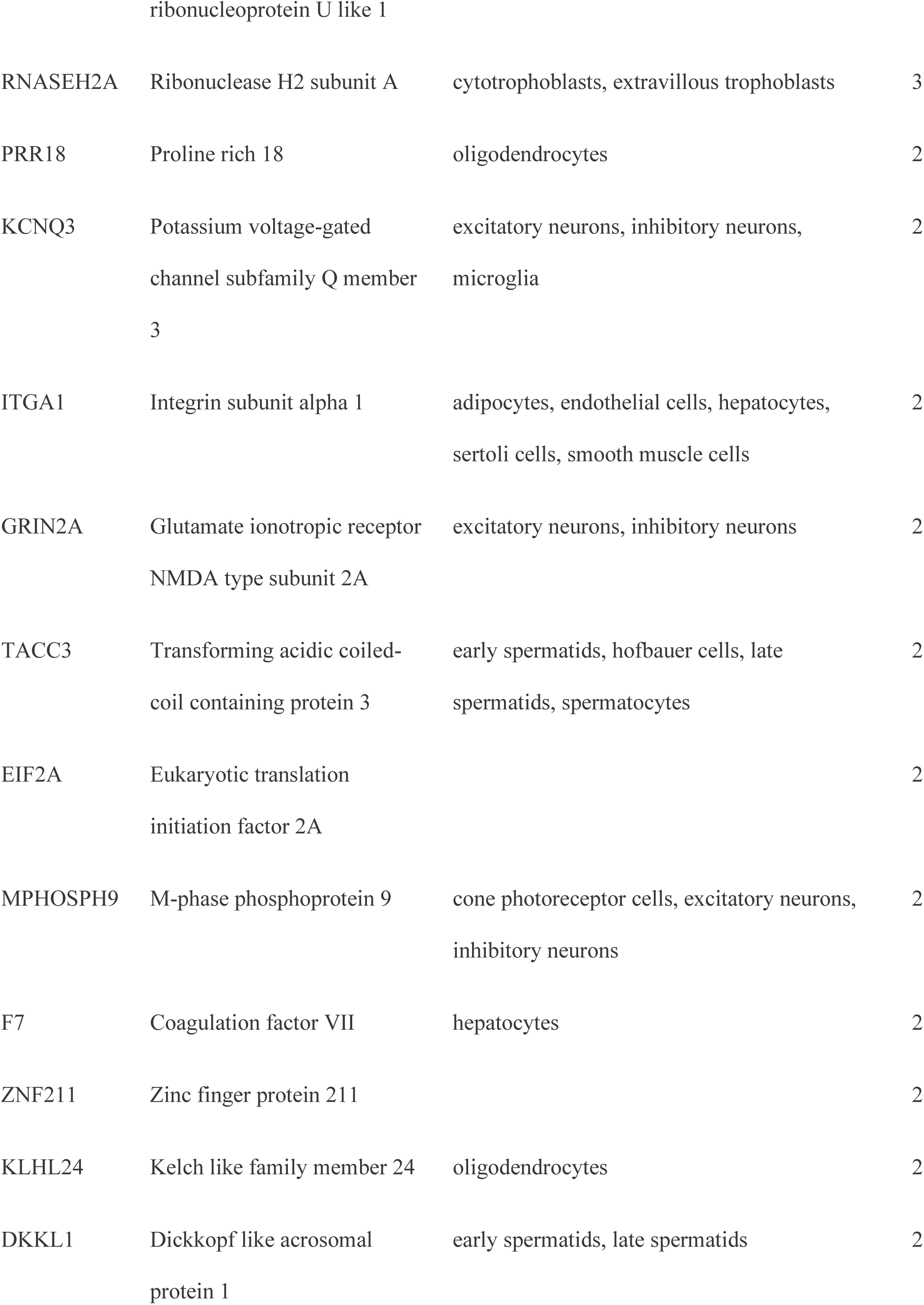
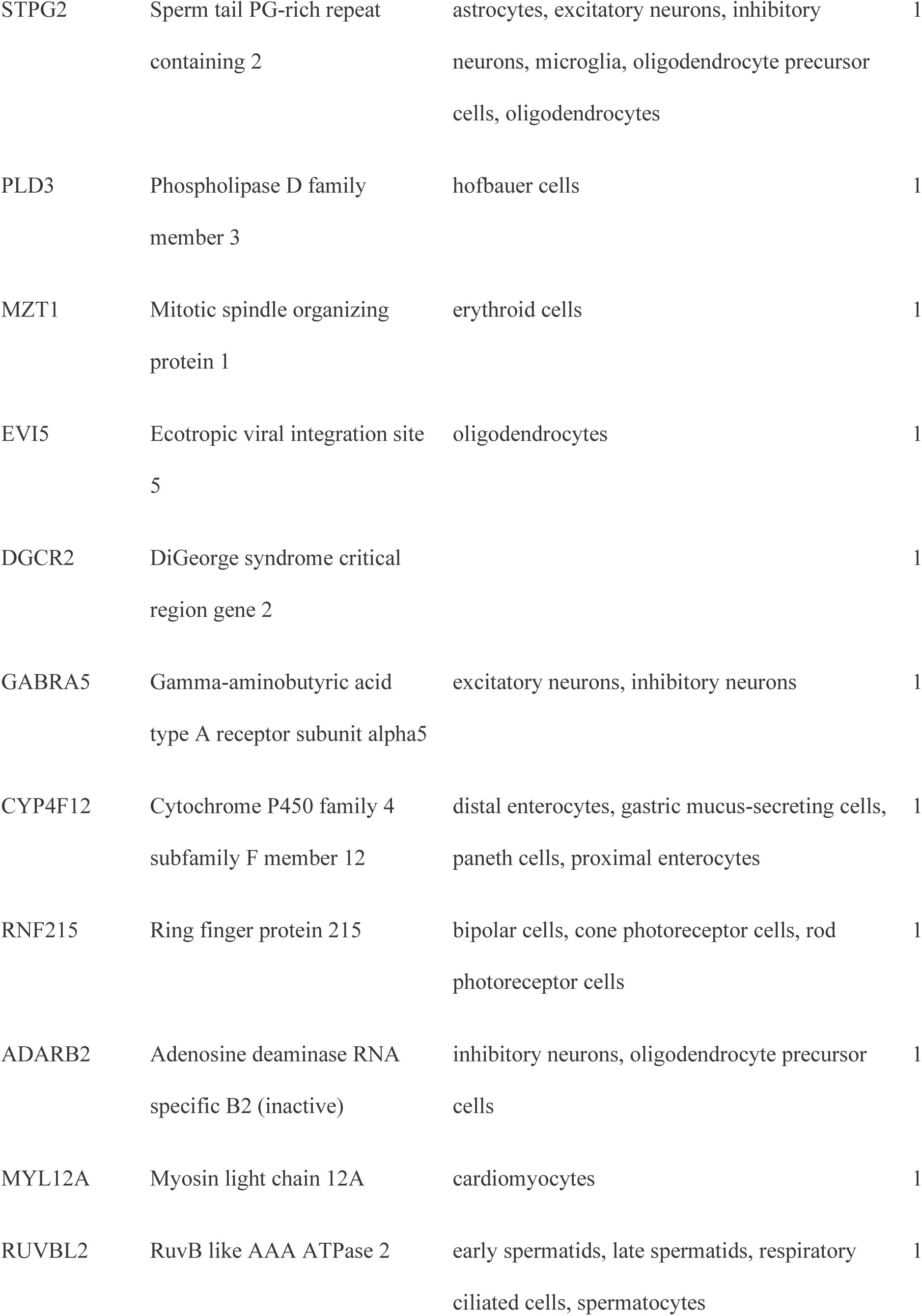
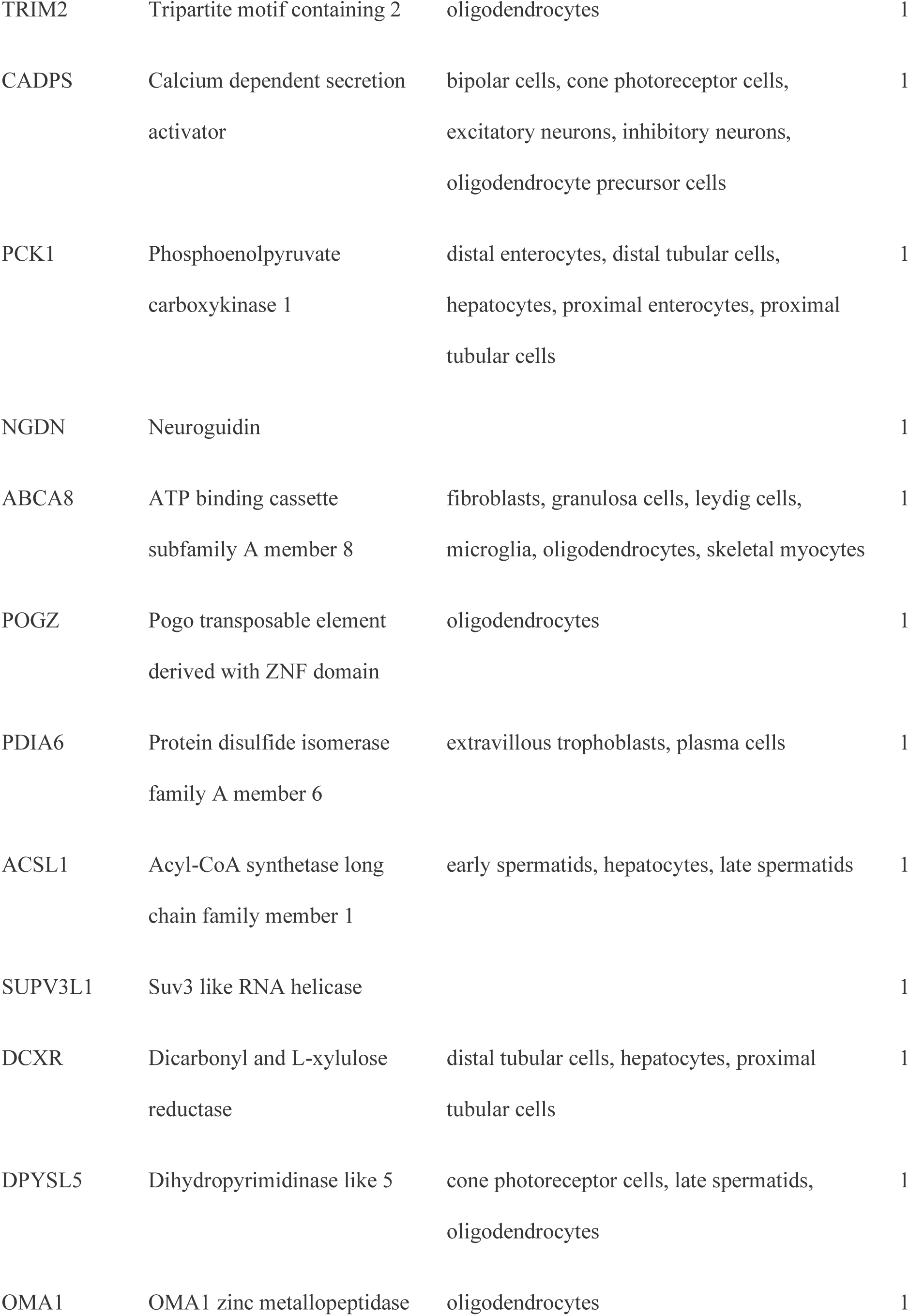
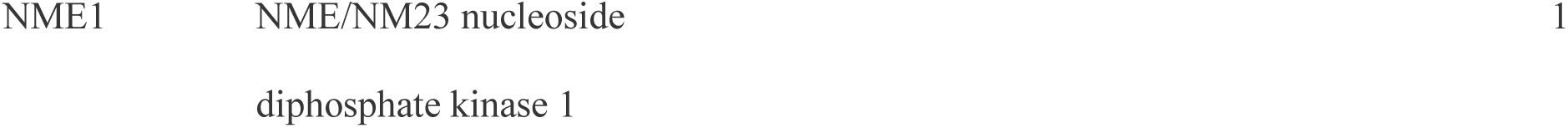
DB TF target genes with differential binding affinity by DB TFs in modern human vs. Neanderthal proximal and semi-distal promoter regions.

The target gene with the greatest number of DB events (143) was FARS2, whose promoter contains 2 SNPs, for which 105 (97.22%) of the DB TF binding models showed differential binding (Table 2). The remaining top ten DB TF target genes with the next highest number of DB occurrences were: TEX19, NARF, SLC6A11, LSM5, FAM172A, WNT7A, EXO1, KRT8, and ZNF37A. Among these genes, all but LSM5 have been associated with brain (FARS2, SLC6A11, FAM172A, WNT7A, and ZNF37A) or reproductive (FARS2, TEX19, NARF, WNT7A, EXO1, KRT8) tissue- or cell-specific expression in the Protein Atlas database (Supplementary Table 2). Among all 74 DB TF target genes, 30 have reproductive-specific expression, 28 have brain-specific expression, 18 have blood cell-specific expression, and 16 have mitochondria-specific expression or localization.

### DB TF target genes and their direct interactors are enriched in the brain, physiology, vision, and autism

The OmniPath DB [26,27] was queried to identify all proteins that have posttranslational interactions with DB TF target proteins. This identified 805 genes that were combined with the 74 DB TF target genes for gene ontology enrichment analysis via the GeneOntology.org webserver [16,17]. Among the most enriched biological process terms associated with physiology were: ‘midbrain morphogenesis’ (GO:1904693; 22x enrichment), ‘positive regulation of connective tissue replacement’ (GO:1905205; 22x), ‘trachea formation’ (GO:0060440; 19x), ‘regulation of midbrain dopaminergic neuron differentiation’ (GO:1904956; 19x), ‘prostate gland morphogenetic growth’ (GO:0060737; 17x), ‘acinar cell differentiation’ (GO:0090425; 17x), ‘vitellogenesis’ (GO:0007296; 17x), ‘frontal suture morphogenesis’ (GO:0060364; 17x), ‘eyelid development in camera-type eye’ (GO:0061029; 15x), and ‘retinal blood vessel morphogenesis’ (GO:0061304; 15x). The complete biological process enrichment results are available in Supplementary Table 3.

Fisher’s exact test revealed enrichment of target genes and their direct interactors among autism spectrum disorder (ASD) genes from the SFARI gene knowledgebase (Banerjee-Basu and Packer 2010) (sfari.org; vers. 3.0), (OR 2.68; p value 1.16 × 10^-16^).

### DB TFs are coexpressed in neural and immune tissues

Data derived from the bulk RNA-Seq experiments (312 different tissues; 686 samples) of the FANTOM5 project were used to generate a cluster map of DB TF expression within the 100 tissues with the highest aggregate expression of the DB TFs, presented in Figure 5A (y-axis DB TFs; x-axis, samples). On the basis of their tissue expression in the FANTOM dataset, the DB TFs form two distinct top-level clusters: the larger cluster contains 89 genes, the majority of which are known to have a role in development (78/89) and specific expression in brain or retina tissue (72/89), whereas the smaller cluster of 17 genes all show specific expression in blood and immune cells. Data on tissue group specificity (brain and retina; male/female) from the Human Protein Atlas (bulk and single-cell RNA-Seq), status as a development-related gene (homeobox, forkhead box, and SRY-related HMG-box), and above the 90th quantile expression in immune cell types as defined by the Database of Immune Cell eQTLs (DICE), were analyzed and included in Figure 5C.

Within the larger cluster of 89 developmental- and brain-specific genes, there is a subcluster of 12 genes whose expression is most limited to FANTOM 5 neural tissues: NKX6-2, CUX2, NR2E1, POU3F2, POU6F2, EMX1, VAX1, POU3F3, DLX1, DLX2, OTX1, and POU3F1. All of these genes are identified in the Protein Atlas as being enriched or enhanced in the brain or retina. Compared with all protein-coding genes, GO analysis of these 12 genes with the PANTHER-based gene ontology.org web server [16,17] revealed the top 10 enriched biological process terms: ‘cerebral cortex GABAergic interneuron fate commitment’ (GO:0021893; 100x enrichment), ’positive regulation of amacrine cell differentiation’ (GO:1902871; 100x), ’forebrain ventricular zone progenitor cell division’ (GO:0021869; 100x), ’negative regulation of photoreceptor cell differentiation’ (GO:0046533; 100x), ’regulation of photoreceptor cell differentiation’ (GO:0046532; 100x), ’regulation of amacrine cell differentiation’ (GO:1902869; 100x), ’neuroblast differentiation’ (GO:0014016; 100x), ’negative regulation of oligodendrocyte differentiation’ (GO:0048715; 100x), ’cerebral cortex GABAergic interneuron differentiation’ (GO:0021892; 100x), and ’negative regulation of glial cell differentiation’ (GO:0045686; 100x).

Additionally, within the brain tissue cluster, we observed subclusters of fetal and newborn tissues (Figure 2D). Overall, DB TFs presented the highest expression in the medial frontal gyrus (newborn), medial temporal gyrus (adult), parietal lobe (fetal), occipital lobe (fetal), eye (fetal), pineal gland (adult), temporal lobe (fetal), retina (adult), occipital cortex (newborn), parietal lobe (newborn), dura mater (adult), medial temporal gyrus (newborn), and spinal cord (fetal) (Supplementary Figure 1). Similarly, the cluster with high expression in blood and immune cells/tissues is comprised of 17 genes: FOS, RORA, FOXK1, FOXK2, IRF3, NFYA, TBP, MAFG, POU2F1, ARID5A, RARA, JUN, CUX1, FOXP1, MEF2C, MAFB, and RXRA. All of these genes have an expression ⋝90th quantile (compared with all protein-coding genes) in at least one immune cell type, as cataloged in the Database of Immune Cell eQTLs (DICE) [28]. GO analysis of the 17 genes (geneontology.org) web server) revealed enrichment of immune-related biological process terms: ‘CD4- positive, alpha-beta T-cell differentiation involved in immune response’ (GO:0002294; 76x enrichment), ‘T-helper cell differentiation’ (GO:0042093; 76x), and ‘alpha-beta T-cell differentiation involved in immune response’ (GO:0002293; 73x), among others.

**Figure 2.**
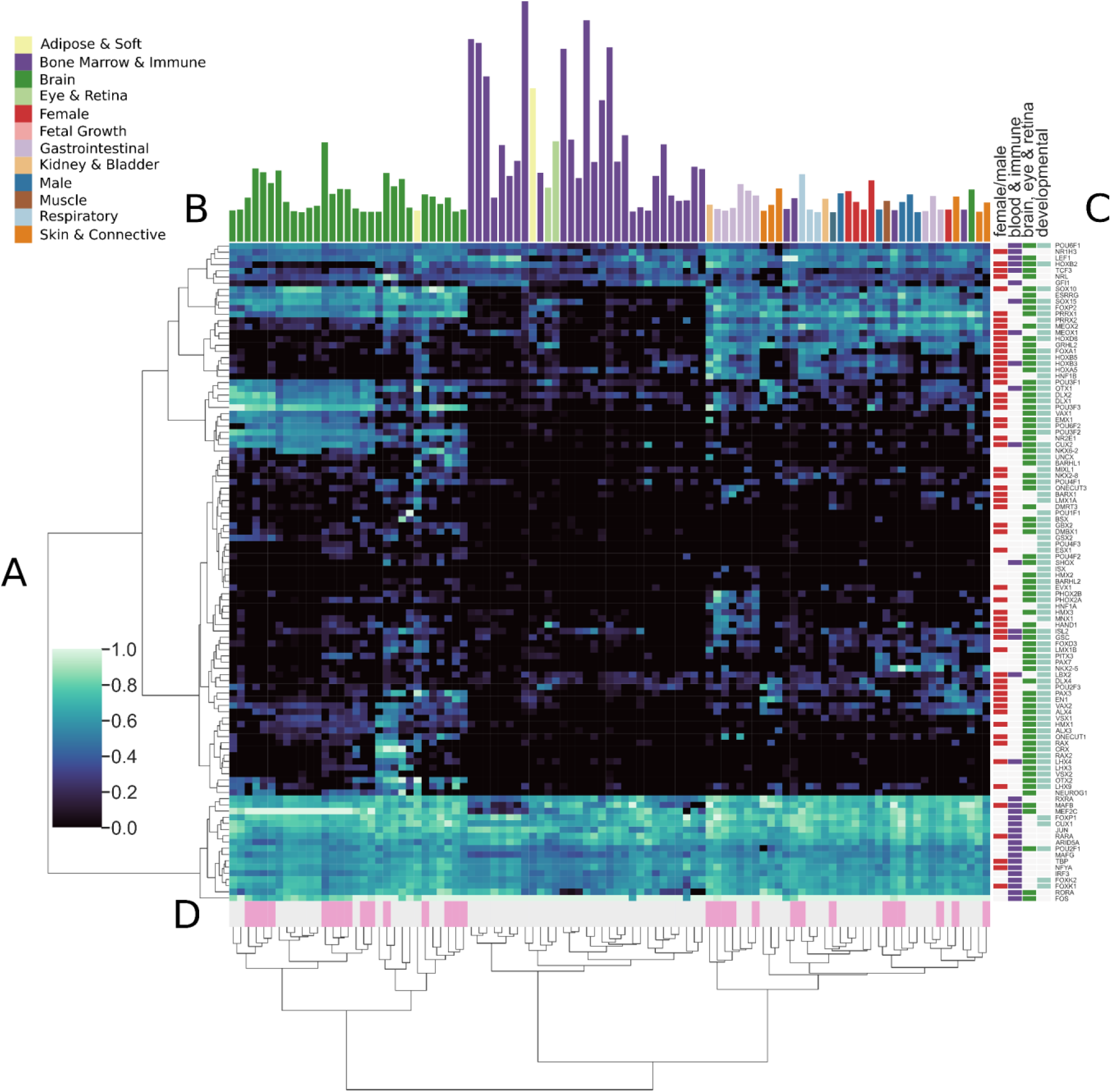
Cluster map of the expression of transcription factors displaying variable binding in modern human vs. Neanderthal. (**A**) Expression data derived from the FANTOM 5 dataset were used to cluster 106 DB transcription factors (y-axis) in 100 tissues (x-axis). Expression data were not available for the DUXA and PROP1 genes. (**B**) Aggregate expression for each tissue across all genes is presented as bars, and bars/tissues are colored by tissue group. Distinct clusters of brain and eye/retina tissues as well as blood and immune cells/tissues are observed. A bar plot with the tissue names is presented in Supplementary Figure 1. (**C**) TF genes are labeled with colored bars corresponding to their identification as homeobox, SOX, or FOX genes (developmental), above the 90th quantile expression in an immune cell type as defined by the DICE database (blood & immune) [28], or to have enriched or enhanced expression by tissue as defined by proteinatlas.org (female/male; brain, eye & retina) [29]. (**D**) Life stage for each tissue is described, where white and pink represent adult and fetal/newborn tissues, respectively. The expression values (TPM) were normalized via log transformation, and each tissue column was adjusted on a zero-to-one scale.

### DB TFs have developmental time-point-related expression profiles in the brain

Cluster analysis of the Allen Brain Atlas bulk RNA-Seq expression data for DB TF genes revealed distinct clusters of time-point-specific expression. Those genes comprising the cluster with the highest aggregate expression in the dataset (module 1 genes: NFYA, TCF3, MEF2C, SOX10, CUX2, TBP, FOXP1, IRF3, SOX15, RARA, FOXK2, NKX6-2, FOXK1, FOS, JUN, POU6F1, POU3F2, MAFG, POU3F3, MAFB, and CUX1) show a dynamic and cyclic pattern of expression during the prebirth timepoints, which then stabilizes in infancy and trends downward in early childhood (Figure 3A). Despite differences in physiology, different tissues generally cluster together by time point rather than tissue type: pcw (8–37 weeks post conception), early (4 months–4 years), and late (8 years–40 years) (Figure 3B).

**Figure 3.**
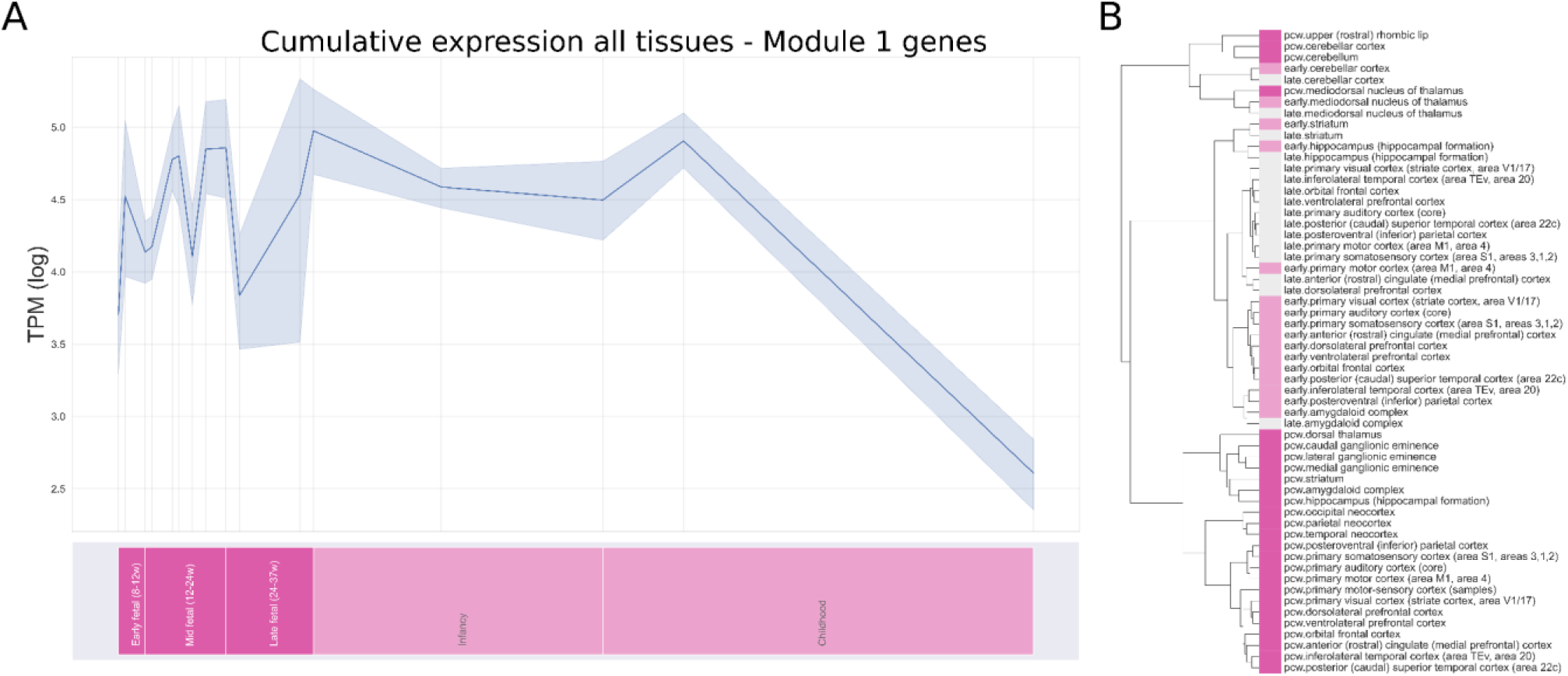
Expression of modern human vs Neanderthal DB TFs in developing brain tissue. The expression data from the Allen Brain Atlas [30] were extracted as RPKM values for 26 unique tissues across 31 timepoints, converted to TPM values, log transformed, and used to construct a cluster map. A) Aggregate expression of the cluster of DB TF genes with the highest total expression across all tissues (module 1) from 8–144 weeks post conception. B) Clustering of brain tissues defined by development time point. The colors represent development stages for both figures: 8–37 weeks post conception (pcw, magenta), 4 months–4 years (early, pink), and 8–40 years (late, white).

### Single-nucleus RNA-Seq of cortical cells reveals differential expression of DB TFs

An analysis of the Allen Brain Atlas single-nucleus RNA-Seq data from 49,417 cortical brain cells [18] was performed to identify which of the MH-Neanderthal DB TF genes may play a functional role in specific annotated brain regions, layers, and cell types. The analyses in the original paper defined metadata indicating cell class (excitatory, inhibitory, and nonneuronal), layer (L1–L6), and predicted type (e.g., astrocyte, microglia, endothelial, etc.), as well as brain regions (Figure 4A). Cell groups defined by the intersection of class, layer, and region were analyzed for differentially expressed genes (DEGs). Several DB TFs were DEGs in specific categories, occurring in nearly all cases in glutamatergic neurons, usually in layers L4–L6, and across nearly all brain regions. The categories in which four or more DB TFs were DEGs are presented in Table 3 and included cut-like homeobox 1 (CUX1), cut-like homeobox 2 (CUX2), estrogen-related receptor gamma (ESRRG), forkhead box protein 1 (FOXP1), forkhead box protein 2 (FOXP2), Myocyte Enhancer Factor 2C (MEF2C), POU Class 6 Homeobox 2 (POU6F2), paired related homeobox 1 (PRRX1), and RAR-related orphan receptor alpha (RORA). In all categories, POU6F2 was a DEG, and in all but two categories, either FOXP2 or FOXP1 (or both) were DEGs. FOXP2 was present as a DEG in excitatory L4 cells in A1C (primary visual cortex), MTG (medial temporal gyrus), S1lm (lower limb somatosensory cortex), and S1 µl (upper limb somatosensory cortex).

**Figure 4.**
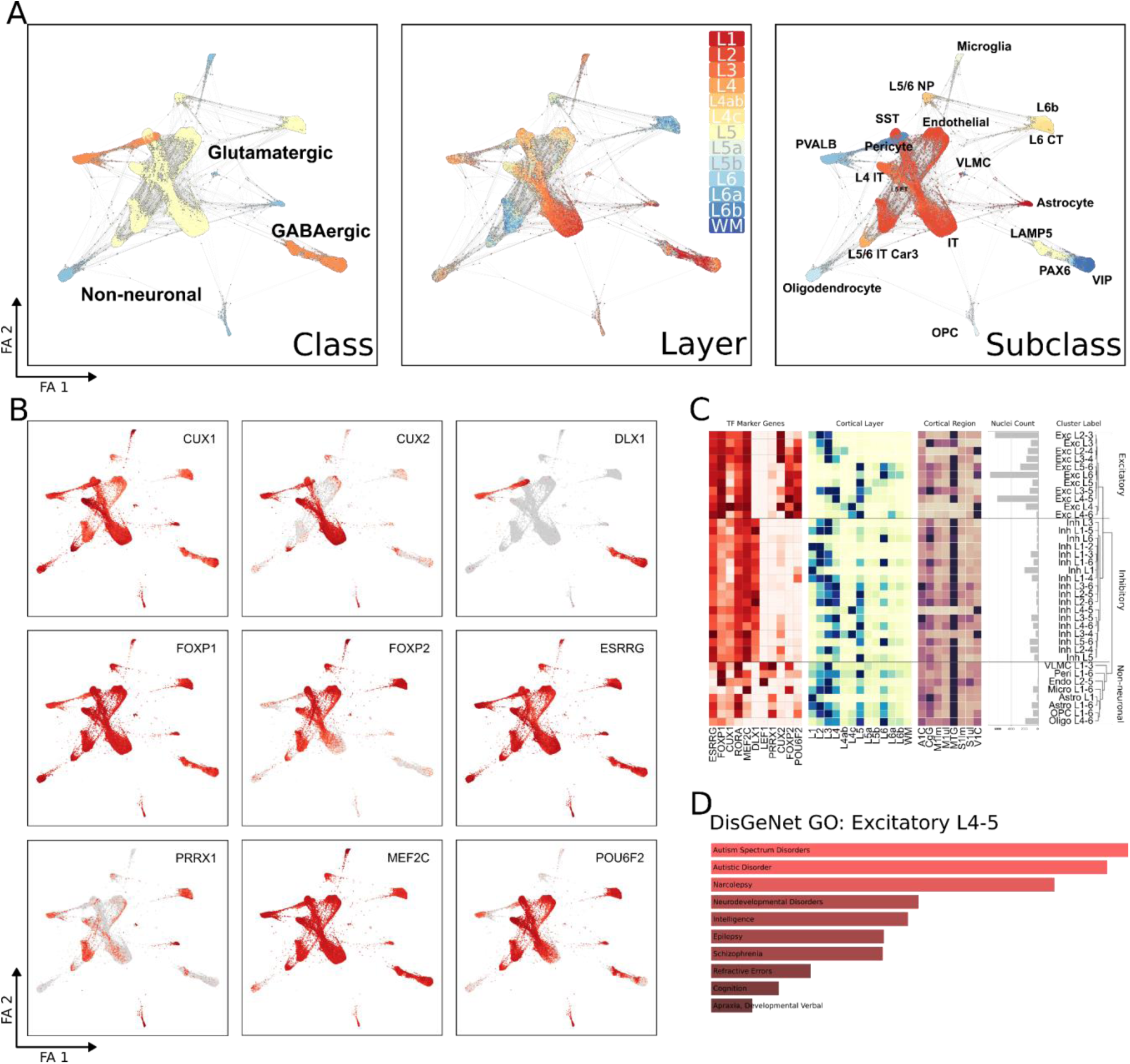
Analysis of single-nucleus RNA-Seq data from brain cortical regions. (**A**) Force-directed graph (FA) visualization of Allen Brain Atlas single-nucleus RNA-Seq data from 49,417 brain cortex cells; the brain excitatory/inhibitory status, cortical layer, and predicted cell type (subclass) data are mapped from annotations in the Allen Brain Atlas sample data [18]. (B) Expression of DB TFs, which are DEGs mapped onto FA visualization. C) On the x-axis: DB TF DEGs, cortical layer, and cortical region; on the y-axis: cell class/type and cortical layer, with the dendrogram indicating expression relationships. The color intensity indicates the normalized expression (0–1 scale). Nuclei count indicates the number of nuclei for that cell class/type at the indicated layer. D) DisGeNET disease ontology terms enriched for the top 100 marker genes for L4–5 excitatory cells. ET, extratelencephalic/pyramidal tract; IT, intratelencephalic; NP, near-projecting; OPC, oligodendrocyte precursor; VLMC, vascular leptomeningeal cell; SST, LAMP5, PAX6, VIP, PVALB, labels on the basis of expression of the corresponding gene.

**Table 3.** DB TFs that co-occur as DEGs in specific regions, layers, and cell types according to the Allen Brain Atlas scRNA-Seq data from the cerebral cortex

| Layer | Class | Region | DEG DB TF genes |
| --- | --- | --- | --- |
| L4 | Glutamatergic | A1C | FOXP2, FOXP1, POU6F2, MEF2C |
| L4 | Glutamatergic | MTG | FOXP2, PRRX1, POU6F2, MEF2C |
| L4 | Glutamatergic | S1lm | FOXP2, PRRX1, POU6F2, ESRRG |
| L4 | Glutamatergic | S1ul | FOXP2, FOXP1, RORA, POU6F2, MEF2C |
| L4 | Glutamatergic |  | FOXP1, CUX2, RORA, POU6F2, CUX1, MEF2C |
| L4 |  | IT | FOXP1, CUX2, RORA, POU6F2, CUX1, MEF2C |
| L4ab | Glutamatergic | V1C | CUX2, RORA, POU6F2, CUX1, MEF2C |
| L4ab |  |  | CUX2, RORA, POU6F2, CUX1 |
| L5 | Glutamatergic | S1lm | FOXP2, PRRX1, POU6F2, ESRRG |
| L5 | Glutamatergic | V1C | FOXP2, FOXP1, RORA, POU6F2, MEF2C |
| L6a |  |  | FOXP2, PRRX1, POU6F2, ESRRG |

### Gene ontology analysis of L4 excitatory neurons

In the Allen Brain Atlas cortical cell snRNA-Seq data, multiple DB TFs were consistently up-regulated DEGs in both layer 4 and excitatory cell groups, and their intersection. Gene ontology analysis of the top 100 marker genes of L4 excitatory cells via the DisGeNet ontology within the Enrichr tool [24] revealed the top enriched terms ‘autism spectrum disorders’, ‘autistic disorder’, ‘narcolepsy’, ‘neurodevelopmental disorders’, ‘intelligence’, ‘epilepsy’, ‘schizophrenia’, ‘refractive errors’, ‘cognition’, and ‘apraxia, developmental verbal’ (Figure 4D). The top 10 enriched biological process terms for L4 excitatory cells were: ‘corticospinal neuron axon guidance’ (GO:0021966; >100x enrichment), ‘corticospinal tract morphogenesis’ (GO:0021957; 91x), ‘ventricular cardiac muscle cell differentiation’ (GO:0055012; 42x), ‘synaptic membrane adhesion’ (GO:0099560; 35x), ’central nervous system projection neuron axonogenesis’ (GO:0021952: 30x), ’synaptic transmission’, GABAergic’ (GO:0051932: 29x), ’positive regulation of excitatory postsynaptic potential’ (GO:2000463: 28x), ’negative regulation of smooth muscle cell migration’ (GO:0014912: 27x), ’positive regulation of dendrite morphogenesis’ (GO:0050775: 25x), and ’modulation of excitatory postsynaptic potential’ (GO:0098815: 25x).

### Gene ontology analysis of L4 excitatory neurons

In the GO analysis of the CC category, the top 10 enriched terms for L4 excitatory cells were: ‘anchored component of presynaptic membrane’ (GO:0099026; 61x enrichment), ‘presynaptic cytosol’ (GO:0099523; 47x), ‘NMDA selective glutamate receptor complex’ (GO:0017146; 47x), ‘intrinsic component of presynaptic active zone membrane’ (GO:0098945; 45x), ‘node of Ranvier’ (GO:0033268; 42x), ‘integral component of presynaptic active zone membrane’ (GO:0099059; 40x), ‘GABA-A receptor complex’ (GO:1902711; 34x), ‘GABA receptor complex’ (GO:1902710; 30x), ‘presynaptic active zone membrane’ (GO:0048787; 27x), and ‘intrinsic component of presynaptic membrane’ (GO:0098889; 22x).

In the GO analysis of molecular function, the top 10 enriched terms for L4 excitatory cells were: ’transmembrane receptor protein tyrosine phosphatase activity’ (GO:0005001; 50x enrichment), ’transmembrane receptor protein phosphatase activity’ (GO:0019198; 50x), ’GABA-gated chloride ion channel activity’ (GO:0022851; 49x), ’inhibitory extracellular ligand-gated ion channel activity’ (GO:0005237; 42x), ’ligand-gated anion channel activity’ (GO:0099095; 35x), ’GABA-A receptor activity’ (GO:0004890; 34x), ’syntaxin-1 binding’ (GO:0017075; 32x), ’GABA receptor activity’ (GO:0016917; 29x), ’transmitter-gated channel activity’ (GO:0022835; 17x), and ’transmitter-gated ion channel activity’ (GO:0022824; 17x).

### DB TFs and Human Accelerated Regions

Doan et al. identified and cataloged a list of genes in close proximity to human accelerated regions (HARs; elevated divergence in humans vs. other primates) as well as those that directly interact with HARs through chromatin conformation changes, e.g., enhancers [31]. Among the set of 110 DB TF genes, 28 were HAR-associated. Fisher’s exact test revealed that the overrepresentation of HAR associations in DB TFs versus all JASPAR TFs was not significant (OR 0.98; p value 0.57). However, compared with all protein-coding genes, the JASPAR TFs as a group were enriched for HAR associations (OR 3.27; p value 6.73 × 10^-26^).

## Discussion

### Regulatory changes drive evolution

Current knowledge indicates that, across the tree of life, prior to and during major radiations of new species, there is a commensurate increase in the number of regulatory genes. Comparative genomic analyses suggest that at the time of eukaryogenesis, or the origin of the last eukaryotic ancestor (LECA), a significant increase in novel TF classes occurred [32]. Eukaryogenesis is one of the major transitions of life on Earth, with increased diversity owing to the endosymbiotic synthesis of an energy-producing α-proteobacteria mitochondrion-progenitor within an archaeon [33]. If the new in-house energy source could be described as the engine driving the diversity arising from LECA, the complementary argument could be that it was TF proteins that performed the steering. Later, prior to the colonization of land by plants, there was an increase in the TF families of the ancestral aquatic streptophytes, which were then already present in the first land plants, which benefited from rapid diversification thereafter in the new environment [34]. Prior to the radiation of bilaterian (multicellular) metazoans, an increase in transcription factor families occurred [35], roughly quadrupling in ratio [36]. Arguably, these increases in TF family numbers immediately prior to major transitions in life could indicate the critical role of TFs in the rise and adaptation of eukaryotes and the complexity arising within their successors.

Far more recently, a transposon-mediated shift in regulatory signaling in mammalian pregnancy has led to the endometrial stromal cell type [37] and the rewiring of a stress response producing the decidual stromal cell type [38]. In humans, the development of our cognitive skills is possibly due to a delay in synaptic-function gene expression and corresponding synaptogenesis in the prefrontal cortex, owing to transcription factors such as myocyte enhancer factor 2A (MEF2A) [39–42]. At the same time, evidence is accumulating that maturation occurred earlier in the closely related Neanderthal, as it does in chimpanzees [43], which is supported by samples of jaw [44], tooth [45], and, most importantly, cranium [46]. Furthermore, the development of a globular brain structure was not present in modern humans at the time of divergence from the Neanderthal and Denisovan lineages but has been a unique product of modern human development within the last 35,000– 100,000 years [47], making it particularly attractive for further regulatory analyses.

### Modern human brain regulatory changes

Compared with the complete set of JASPAR TFs, which are very strongly enriched compared with all protein-coding genes, the TF genes identified as differentially binding genes are strongly enriched for homeobox genes, the primary regulators of development in humans and other multicellular organisms. Similarly, the DB TFs are enriched for brain-specific expression compared with the JASPAR TFs, which are again enriched compared with all human genes. Hierarchical clustering of TF expression revealed that DB TF genes associated with neural and immune functions formed distinct groups. Within the tissue axis, in the neural cluster, adult and fetal/newborn neural tissues cluster separately in several instances (Figure 2). From these discoveries in a broader set of data, it became important to focus more closely on the expression of these DB TFs in developing neural tissues specifically. Analysis of RNA-Seq data from whole-brain subtissues further revealed differences in the expression of these genes across time and likely in a developmentally dependent manner. Specifically, the expression of DB TFs in all tissues for the two groups occurring within 8 weeks post conception to 4 years post birth was largely segregated from all tissues for the group defined by 8 years to 40 years of age (Figure 3). Additionally, several modules of DB TFs identified by cluster analysis appear to show a cyclical pattern of expression during fetal development, which then stabilizes post birth (Figure 3A).

### DB TFs co-occur as DEGs according to single-cell RNA-Seq of glutamatergic cells in the L4 human cortex

Recently, the proliferation of single-cell transcriptomics has begun to clarify the direct role of transcription factors in cell type determination and identity (Arendt et al. 2019). To determine in which cell types DB TFs are expressed in the adult brain, single-nucleus data for 49,417 cortical cells were examined. A subset of the DB TFs were annotated as commonly co-occurring DEGs: CUX1, CUX2, ESRRG, FOXP1, FOXP2, MEF2C, POU6F2, PRRX1, and RORA. Interestingly, we observed that these DB TF DEGs were primarily co-occurring in glutamatergic cells of the L4 cortex. Among this set of nine genes, six (CUX1, CUX2, FOXP1, FOXP2, MEF2C, and RORA) are cataloged in the SFARI Gene knowledgebase [48] (sfari.org; vers. 3.0) as ASD candidate genes. Similarly, six (CUX1, FOXP1, FOXP2, MEF2C, POU6F2, and RORA) are present in the genome-wide association studies (GWAS) catalog [49] (www.ebi.ac.uk/gwas) for the ‘schizophrenia’ (MONDO_0005090) term. In support of this connection, a recent large-scale study on schizophrenia genetics in 74,776 individuals with schizophrenia (and 101,023 control individuals) revealed enrichment of genes that are highly expressed in glutamatergic neurons of both the human and mouse cerebral cortex [50]. Similarly, a recent large-scale study on ASD genetics in 11,986 individuals with ASD (and 23,598 control individuals) identified 102 ASD risk genes and revealed that fetal-development excitatory neurons accounted for the greatest number of these genes among all the identified cell types [51].

### Co-occurring DB TF DEGs associate with HARs and neuropsychiatric disease

Human-accelerated regions (HARs) are locations in the genome where the rate of evolutionary change has accelerated since divergence from chimpanzee. A study of 2,737 HARs revealed that they are enriched for TFBSs generally and for TFs associated with neural development specifically (Doan et al. 2016). While the DB TFs were not enriched for HAR association compared with the JASPAR TFs (OR 0.98; p value 0.57), the JASPAR TFs are compared to all genes (OR 3.27; p value 6.73 × 10^-26^). Using the HARs cataloged by [31], we identified 28 DB TFs that are either in close proximity to a HAR or directly interact with one through chromatin conformation changes. Additionally, all but one (CUX2) of the co-occurring DEGs DB TFs (CUX1, ESRRG, FOXP1, FOXP2, MEF2C, POU6F2, PRRX1, and RORA) identified in the L4 excitatory neurons of the cerebral cortex (Allen Brain Atlas scRNA-Seq data) contain HAR regions or HAR interactivity. Several of these contain or associate with HAR variants that have been noted to cause neurological disease. POU6F2 possesses an intronic HAR in which a mutant allele (GRCH38:chr7:39,033,595) is associated with ASD (Doan et al. 2016). The promoter of the CUX1 gene has been determined via ChIA-Pet analysis of chromatin to interact with a HAR located ∼200 kb away; unrelated individuals with intellectual disability (ID) (IQ<40) and ASD have been identified with a homozygous mutation (GRCh38:chr7:101,606,361) in this HAR. A luciferase reporter assay revealed that when the mutant version of this HAR interacts with the promoter of the CUX1 gene, its expression is increased threefold, while cultured differentiated neurons with increased CUX1 expression exhibit increased synaptic spine density. Similarly, another HAR (GRCh38:chr5:88,480,873) has been shown to interact with the promoter of MEF2C, and a mutation in this HAR creates a putative MEF2A binding site, reducing expression by ∼50%. Mutations in the MEF2C gene are associated with autism [52,53], mental retardation [52–54], schizophrenia [55,56], epilepsy [52,53], and speech abnormalities [52]. Additionally, binding motifs for several of the DB TFs identified in our study were found to be enriched in HAR regions; POU6F1 and POU2F1 in ultra-conserved HARs, and HNF1A in all HARs [31]. After these intriguing results, further direct analysis of HARs via the TFBSFootprinter tool is warranted.

Importantly, one of the identified DB TFs in our data is the FOXP2 TF gene, which was the first noted "speech gene" [57]. For the early postnatal period in mice, FOXP2 is a negative regulator of MEF2C, and the likely result is the promotion of synaptogenesis in the cortical striatum [58]. In addition to this interaction with MEF2C, there is a POU3F2 (which, according to our results, is also a DB TF) binding site within the FOXP2 gene, which has been shown to affect FOXP2 expression and is associated with a selective sweep that has occurred since the divergence of humans and Neanderthal [59,60]. Because of its significant role in human evolution, further analysis is warranted regarding the regulation of FOXP2 in other relevant areas (e.g., introns, 3’ UTRs, enhancers, etc.) and for genome-wide binding sites of FOXP2 itself.

### DB TF target genes are expressed in oligodendrocytes, testes/sperm, the maternal–fetal interface, and mitochondria

There were 74 genes identified as DB TF target genes. All but 8 of these genes are specifically expressed in reproductive (30), brain and retina (28), immune (18), or mitochondrial (16) cells or tissues. Among the 30 DB TF target genes with reproductive cell- and tissue-specific expression, the majority (19) are expressed in male tissues (testis, early and late spermatids, spermatocytes, and spermatogonia). A total of 12 genes were expressed in female-specific cells and tissues and, in many cases, specifically in those involved in the maternal–fetal interface (cervix, granulosa cells, cytotrophoblasts, extravillous trophoblasts, syncytiotrophoblasts, and endometrial ciliated cells). Among the 28 DB TF target genes with brain- and retina-specific expression, 16 are expressed in oligodendrocytes, 12 in inhibitory neurons, and 11 in excitatory neurons.

The DB TF target gene with the greatest number of differential binding events in its promoter region is FARS2, a member of the mitochondrial aminoacyl-tRNA synthetases (mt-aaRSs). scRNA-Seq data from the Protein Atlas indicate that FARS2 expression is increased in excitatory neurons, oligodendrocyte precursor cells, inhibitory neurons, oligodendrocytes, astrocytes, microglia, late spermatids, and early spermatids. mt-aaRSs charge their cognate mt-tRNA with the appropriate amino acid, and mutations in these genes cause a variety of diseases, most predominantly those that affect the central nervous system [61], perhaps due to delayed myelination, demyelination, or both [62]. In addition to their canonical roles, mt-aaRSs are hypothesized to function in monitoring amino acid levels, as sensors for the mitochondrial environment, and in transcriptional regulation [61], and through the addition of new protein domains have been associated with neural development and immune response, among other processes, as reviewed by [63]. The FARS2 gene produces the mt-PheRS protein (phenylalanyl-tRNA synthetase), which is mitochondria-locating and is responsible for attaching phenylalanine to its corresponding mt-tRNA for mitochondrial protein translation [64]. Intragenic variants in the FARS2 gene have been linked to two primary clinical manifestations, early-onset epileptic mitochondrial encephalopathy and spastic paraplegia, and for patients in both groups, symptoms can also include intellectual disability or developmental delay [64]. Deletions within FARS2 and reduced expression levels have also been associated with schizophrenia [65].

Importantly, FARS2 shares a bidirectional promoter with another mitochondrial gene, LYRM4 (LYR motif-containing protein 4), which has a TSS just ∼400 bp away. Together with ISCU, NFS1, and FXN, the ISD11 protein of the LYRM4 gene is involved in the formation of iron–sulfur (Fe–S) clusters [66,67], which are essential cofactors in many basic biological processes, such as the formation of respiratory chain complexes I, II, and III [68] and subsequent oxidative phosphorylation [69]. Mutations in LYRM4 cause deficits in oxidation phosphorylation reactions [70], which are critical in neuronal development and schizophrenia, as reviewed previously [71]. In support of this connection, polymorphisms occurring in the FARS2-LYRM4 bidirectional promoter region have been shown to be associated with cognitive deficit and schizophrenia [72]. Similarly, a 900 kb microdeletion in this region (encompassing RPP40, PPP1R3G, LYRM4, and part of the FARS2 and CDYL genes) has been shown to produce gyral pattern anomalies, intellectual disability and speech and language disorders [73].

A comparison of metabolite levels in the prefrontal cortex, visual cortex, cerebellum, kidney, and muscle between humans and other primates (chimpanzee and macaque), showed ‘aminoacyl-tRNA biosynthesis’ was the most significantly enriched pathway for metabolites having higher levels in human; for all three brain regions, but not muscle or kidney [74]. Similarly, the ‘Phenylalanine, tyrosine and tryptophan biosynthesis’ pathway was enriched for all three brain regions. This observation fits with the ongoing hypothesis that there is a requirement for the coevolution of nuclear and mitochondrial genes coding for mitochondrial proteins, known as mitonuclear compensatory coevolution, as reviewed in [75], and that this coevolution is a driver of speciation [76,77]. In the mitonuclear compensatory coevolution hypothesis, nuclear-encoded genes encoding aminoacyl tRNA synthetases (such as FARS2) and OXPHOS complex components (such as LYRM4) are expected and, in some cases, have been shown to evolve more rapidly in response to high rates of evolutionary change in mitochondrially encoded genes, with which their protein products subsequently directly interact [75]. Another consideration is how to regulate the translation of OXPHOS complex components when the corresponding genes exist in both the mitochondrial and nuclear genomes [78,79]. The high number of observed DB TFBSs in the bidirectional promoter of the nuclear FARS2 and LYRM4 genes thus potentially allows more well-regulated coordination of cross-compartment OXPHOS component gene expression and may even contribute to the effects of both mitonuclear compensation and speciation.

Taken together, the LYRM4-FARS2 locus is potentially of great interest in the divergence of modern humans from other hominins and hominids. It is reasonable to imagine that differences in neuronal development and function, and even diet, between MH and Neanderthal could require corresponding differences in mitochondrial activity, metabolism, and amino acid sensing.

### Lots of work left to do

The divergence of modern humans and Neanderthals is estimated to have occurred between 400,000 and 800,000 years ago [80–82]. Since then, both subspecies have experienced unique evolutionary, social, and cultural paths, many of which overlap and intertwine. What appears to be both divergent and unifying in the hominin lineage is a significant change in the brain, leading to differing abilities in terms of cognition, social function, language and creativity. In the case of humans, and likely for Neanderthals as well, many of the same genes, which are at the core of our novel capabilities, are the same which are commonly the root of our maladies. Among other neuropsychiatric disorders, autism and schizophrenia appear to be maintained in humans at a greater rate than would be expected if they were strictly deleterious: the global prevalence of ASD is approximately 1% [83], and 0.28% for schizophrenia [84]. Likewise, as the study of these diseases continues, the direction of understanding tends toward a continuum of effects rather than a discrete on/off state. A study by Linscott et al. reported that the prevalence of psychotic experiences in the general population was 7.2% [85]. Our results revealed that DB TFs are enriched for development and the brain, and studies on HAR regions have shown that mutations in regulatory regions alone are sufficient to cause significant neuropsychiatric effects [31]. As a result, there appear to be significant opportunities to analyze gene regulation to understand not only the evolution of human cognitive abilities but also the neuropsychiatric disorders that accompany them.

### Limitations

We have not included analyses of 3’ UTRs, introns, or enhancers. The single-most expressed transcript for each protein-coding gene was chosen for analysis, while MH vs. Neanderthal SNPs may have occurred more closely to TSSs of lower-expressed transcripts and thereby had higher and perhaps more significant scoring. Only transcripts for protein-coding genes were analyzed, despite the existence of numerous other transcribed sequence types. The SNP dataset we used for analysis is based on multiple Neanderthal individuals and has lower sequencing coverage than newer datasets. The inclusion of multiple individuals is useful for comparing modern humans to Neanderthals as a group but can provide only a more general comparison. Using a higher coverage dataset would allow greater assurance that the SNPs under inquiry are legitimate.

## Acknowledgments

The nonprofit CSC—IT Center for Science Ltd., owned by the state of Finland and Finnish higher education institutions—is acknowledged for providing computational resources for analyses.

## Data availability

Code reproducing the analyses and figures in the manuscript are available on GitHub at https://github.com/thirtysix/TFBS_footprinting_manuscript. The results of the MH vs. Neanderthal promoter analysis are available at https://osf.io/r2mtw/.

## Funding

This work was supported by the Finnish Cultural Foundation and Fimlab to HB and the Academy of Finland and the Jane & Aatos Erkko Foundation to SP.

## Conflicts of interest

The authors declare that they have no conflicts of interest.

---

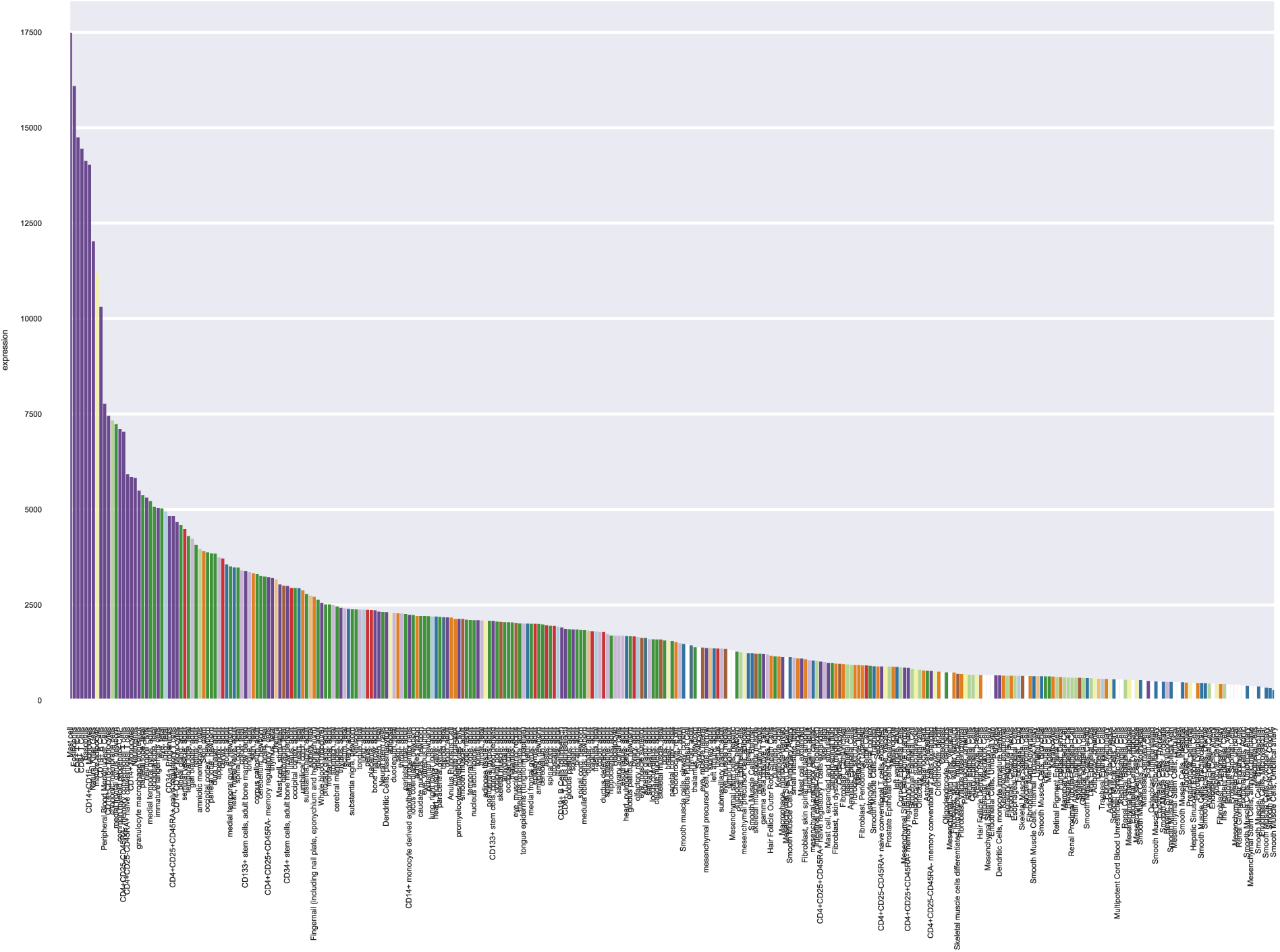

## Notes

https://github.com/thirtysix/TFBS_footprinting_manuscript

https://osf.io/r2mtw/

### Competing Interest Statement

The authors have declared no competing interest.

### Summary of Updates

Part of the methodology (explanation of TFBSFootprinter tool) has been removed, which is becoming part of a new article. This significantly modified the text (primarily removal). General language revisions.

